# A fully human pluripotent stem cell-derived blood-brain barrier model for clinically relevant disease modeling and selection of neurotropic adeno-associated viruses

**DOI:** 10.64898/2025.12.21.695793

**Authors:** Angelo Iannielli, Serena Gea Giannelli, Thomas Behrens, Mirko Luoni, Sharon Muggeo, Giosuè Moscato, Martina Greco, Elisa Enzi, Rita El Khoury, Elena Criscuolo, Monica Casucci, Nicola Clementi, Josep M Canals, Raffaele Iorio, Vania Broccoli

**Affiliations:** CNR-Institute of Neuroscience, 20129 Milan, Italy; Stem Cell and Neurogenesis Unit, Division of Neuroscience, IRCCS San Raffaele Scientific Institute, 20132 Milan, Italy; Innovative Immunotherapies Unit, Division of Immunology, Transplantation and Infectious Diseases, IRCCS San Raffaele Scientific Institute, 20132 Milan, Italy; Laboratory of Microbiology and Virology, Vita-Salute San Raffaele University, Milan, Italy; Laboratory of Stem Cells and Regenerative Medicine, Department of Biomedical Sciences, Faculty of Medicine and Health Sciences, University of Barcelona, 08036 Barcelona, Spain; Creatio, Production and Validation Center of Advanced Therapies, Faculty of Medicine and Health Sciences, University of Barcelona, 08036 Barcelona, Spain; August Pi i Sunyer Biomedical Research Institute (IDIBAPS), 08036 Barcelona, Spain; Dipartimento Universitario di Neuroscienze, Università Cattolica del Sacro Cuore, Fondazione Policlinico Universitario Agostino Gemelli IRCCS, Rome, Italy

## Abstract

The blood-brain barrier (BBB) is a highly functionalized vascular interface which regulates the exchange of substances between the neural parenchyma and its periphery. BBB leakage, leading to its uncontrolled permeability, is increasingly recognized to facilitate the onset of neuropathologies and aggravate their clinical progression. In vitro models of the BBB have rapidly evolved into elaborated structures that mimic its spatial architecture and multicellular nature. However, their cellular components are currently highly heterogeneous in origin and maturation state. Here, we have developed novel procedures to establish reproducible and scalable sources of endothelial, mural and astroglial cells generating a fully human pluripotent stem cell (hPSC)-derived BBB model, termed thBBBA. hPSC-derived BBB cell types are readily assembled into thBBBAs that develop mature functional properties with high barrier impermeability. Mature thBBBAs can also be generated by frozen hPSC-derived cell samples, providing a simple and scalable off-the-shelf system for general use. thBBBAs were instrumental in identifying the critical pathological role of an IL-6 autocrine source in disrupting thBBBA integrity, increasing its permeability to NMDAR antibodies from autoimmune encephalitis patients, and revealing the therapeutic effects of tocilizumab in this setting. Additionally, we have shown that thBBBAs are an invaluable system for ranking the clinical readiness of novel engineered AAV neurotropic capsids, previously selected in animal models or in vitro systems.

## Introduction

The brain vasculature is distributed throughout the neural tissue in a pervasive manner, reaching proximity to the majority of neurons, that have a high energy demand and require a continuous supply of nutrients and oxygen. The brain vessels are organized in a three-cellular layer structure, called the blood-brain barrier (BBB), which restricts the transport of compounds, noxious agents and cells from the blood to safeguard the fragile neural environment (*1,2*). BBB is constituted by endothelial cells (ECs) lining the blood vessels, mural cells (MCs) in close contact with the ECs, and astroglial cells (ACs) extending their end-feet to the abluminal side of the vessels. This configuration leads to the formation of robust tight junctions between ECs, which minimize paracellular transport. In addition, transcellular transport is also reduced due to a decrease in transcytosis vesicle trafficking (3,4). This high impermeability of the BBB poses a formidable obstacle to the delivery of therapeutics into the brain to exert their therapeutic effects in the treatment of neurological disorders (*5*). In vitro models of the BBB are pivotal to our understanding of the physiological mechanisms that modulate its functional properties in health and disease, and to predicting the crossing potential of novel compounds (*6,7*). Human multicellular models have recently been generated to reconstitute the full cellular composition of the BBB by mixing different combinations of primary, immortalized or hPSC-derived cells. For example, endothelial cells in these assemblies are commonly obtained from immortalized stable cell lines (e.g. CMEC/D3, HBEC-5i) or directed differentiation of hPSCs. Although stable cell lines provide a rapidly expandable source, these endothelial cells fail to achieve full functional maturation, producing endodermal layers with low tightness and impermeability (*8*). A benchmark protocol for the differentiation of hPSCs into brain microvascular endothelial cells (iBMECs) is based on the single-step induction of a mixed population of neural and EC progenitors (ECPs). ECPs are then expanded in an endothelial basal medium and their differentiation is induced by adhesion to collagen IV/fibronectin-coated plates with a cocktail of defined supplements (*9,10*). Despite forming a monolayer with high barrier integrity, hPSC-derived iBMECs exhibit low expression of some canonical endothelial markers and upregulation of epithelial-related genes, as revealed by transcriptional profiling (*11*). Thus, these conflicting results may indicate that hPSC-derived iBMECs have not reached a fully defined cell specification and technical improvements are needed to generate a more reproducible source of human brain ECs. Similarly, it is not well defined for how long primary mural cells or astrocytes directly isolated from human brain tissue maintain unaltered their phenotype in culture. Therefore, generating high quality cellular sources is a fundamental requisite to obtain BBB models with improved reproducibility and fidelity displaying functional properties as similar as those of its authentic in vivo counterpart. Herein, we set out to generate the three BBB cell types by optimizing hPSC differentiation protocols that combine small molecules and the expression of cell lineage-determining transcription factors (TFs). These advanced procedures have led to the generation of a BBB model reconstituted solely from hPSC-derived cells, with mature functional properties that confers high barrier strength and impermeability. This BBB platform has proven invaluable in defining novel neuropathological mechanisms triggered by autoimmune encephalitis and evaluating the translational potential of novel engineered AAV vectors.

## Results

### hESC differentiation into the three different BBB cell types with high efficiency and fidelity using cell lineage-determining transcription factors

With the aim of establishing a fully hESC-derived BBB structure, we set out to develop efficient differentiation protocols that could facilitate the generation of its three cellular components with consolidated functional maturity. Endothelial cells were differentiated using a well-established procedure based on simultaneous TGFβ inhibition and supplementation with angiogenic factors to enhance vascular cell commitment and proliferation (Fig. 1a) (*11,12*). However, non-endothelial cells negative for the key marker PECAM1 were found to be a common contaminant in these cultures, complicating their use for this application (Fig. 1b). We, therefore, complemented the small-molecule cocktail with the lentiviral transduction of the two ETS factors ETV2 and ERG1 in various combinations. These TFs are key determinants of the vascular development and have been used as genetic tools for reprogramming non-vascular cells (*13,14*). Lentiviral-based overexpression of ETV2 or ERG1 alone promoted a more homogeneous endothelial differentiation, but the cell monolayer resulted disorganized with sketchy expression of cell junction markers (Supplementary Fig. 1). In contrast, when both factors were co-expressed together the majority of the differentiating hESCs into PECAM1+ endothelial cells (ECs) with a regular cobblestone-like morphology (Fig. 1b; Supplementary Fig. 1). Thus, we generated a hESC line (EE-hESC) stably integrating the two lentiviral vectors at low copy numbers, as evaluated by qPCR vector quantification assays, for doxycycline (dox)-inducible expression of ETV2 and ERG1 (2.3 and 3.1 copies in average per cell for the ETV2 and ERG1 vectors, respectively) (Fig. 1c). EE-hESC lines maintained unaltered the pluripotent stem cell state as suggested by the constitutive expression of key markers and exhibited a normal karyotype (Supplementary Fig. 2a-c). Nearly all EE-hESCs, exposed to the same small molecule cocktail along with dox for 2 weeks, differentiated into PECAM1+ ECs at DIV25 (99.6 ± 0.2%). At confluence, EE-ECs acquired a cobblestone morphology with well-organized cell junctions enriched in OCCLUDIN, GLUT1 and ZO-1, forming a regular epithelial monolayer (Fig. 1d). We next used RNA sequencing (RNA-seq) to profile global gene expression in DIV30 hESC-derived ECs obtained with the small molecule cocktail with or without ETV2/ERG1 (EE-ECs and SM-ECs, respectively) and CMEC/D3 brain microvascular endothelial cells, which are widely employed as the endothelial component of in vitro BBB structures (*6,15*). Overall, the transcriptional profile of EE-ECs appeared to be highly divergent from that of SM-ECs and CMEC/D3 as revealed by both principal component analysis (Fig. 1e) and unsupervised hierarchical clustering (Fig. 1f) of the whole transcriptomes. Analysis of transcripts in EE-ECs compared to SM-ECs and CMEC/D3 cells revealed several enriched Gene Ontology (GO) terms, including vascular development and differentiation and cell adhesion regulation, while terms associated with mesenchymal cell fate and epithelial-to-mesenchymal transition were downregulated (Supplementary Fig. 3a,b), indicating that vascular cell identity was better represented in EE-ECs. Consistently, EE-ECs show the highest correlation with primary human brain endothelial cells in terms of global gene expression profiling in a pairwise comparative analysis (Supplementary Fig. 3c) (*16*). In particular, cardinal genes of vascular cell specification and function such as CDH5, ENG, ERG and ICAM1/2 were enriched in EE-ECs, whereas SM-ECs and CMEC/D3 cells displayed high expression of distinct mesenchymal cell markers such as SNAI2, COL1A1/2 and NID2 (Fig. 1g). In contrast, CMEC/D3 cells clustered very apart from both types of hESC-derived ECs and displayed high expression levels of many mesenchymal-associated genes, suggesting that their endothelial identity had been eroded during their in vitro expansion. Furthermore, EE-ECs exhibited negligible expression of major epithelial-specific genes such as SOX9, EPCAM, CDH1, GRHL2, OVOL1, ELF3, and MUC1, thereby ruling out a differentiation bias toward the epithelial lineage (Fig. 1g). These findings indicate that, under this culture condition, EE-ECs established a robust endothelial phenotype and upregulated key BBB-specific markers. We though that this could be due to the presence of CHIR99021 in the differentiation small-molecule cocktail, a strong activator of the Wnt pathway, which is known to play a pivotal role in promoting the BBB endothelial identity during development (17,18). In fact, its removal from the differentiation mixture strongly impaired the endothelial cell maturation and blunted the expression of the BBB endothelial-specific genes (Supplementary Fig. 4a,b). Altogether, these data confirm that the combination of small molecules with expression of the master regulators ETV2/ERG1 is superior for differentiating vascular cells with high efficiency and fidelity.

**Figure 1:**
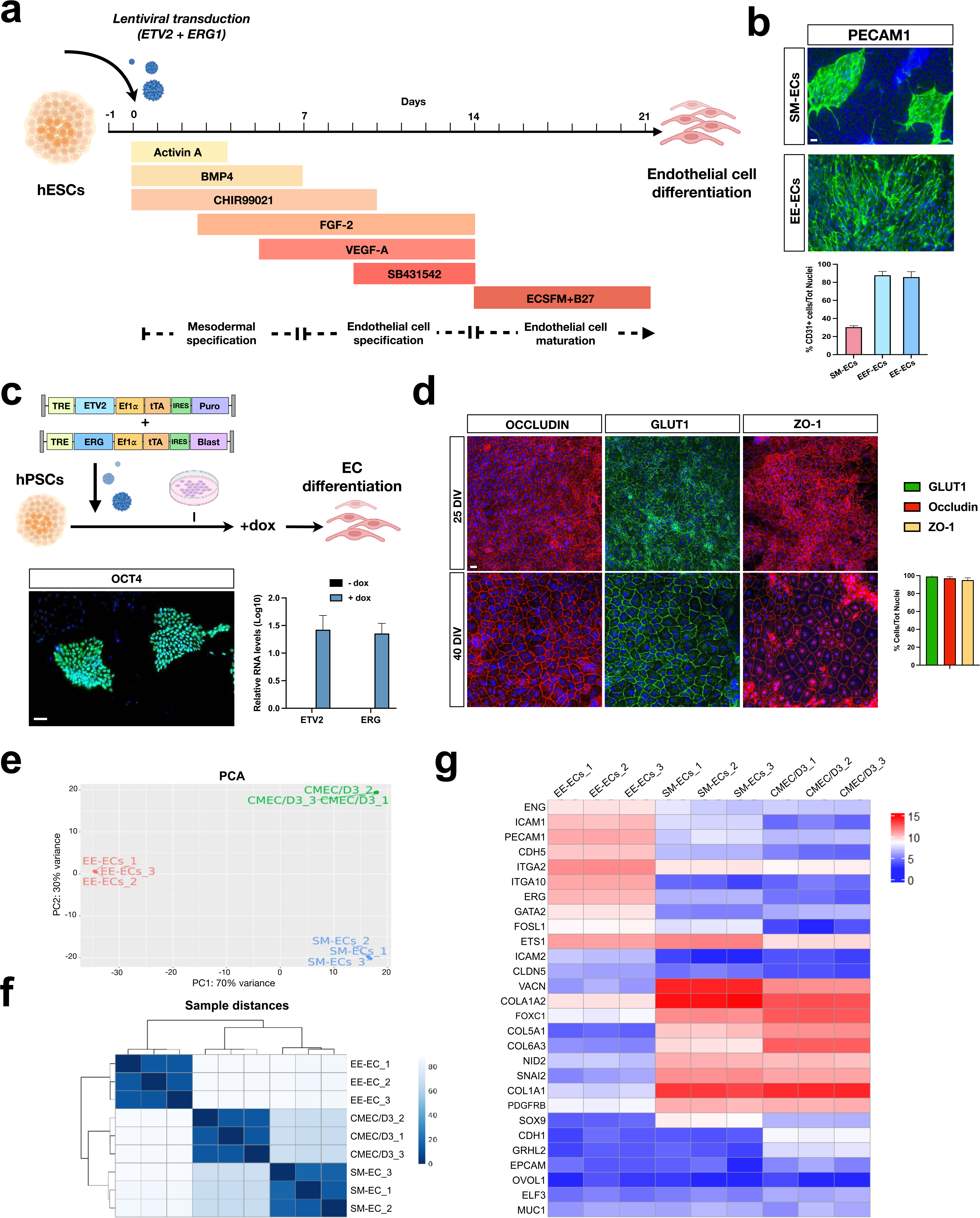
Generation of hESC-derived endothelial cells. (**a**) Illustration of the procedure for differentiating endothelial cells from hESCs. (**b**) Representative images and quantification of hESC-endothelial cells immunostained for PECAM1 (green) (n=3 independent experiments). (**c**) Diagram of the protocol for generating ETV2 and ERG stable cell lines (EE-hESC). Representative images of EE-hESC colonies and qPCR analysis of ETV2 and ERG1 expression with or without doxycycline (n=3 independent experiments). (**d**) Representative images and quantification of hESC-derived endothelial cells immunostained for the specific markers OCCLUDIN, GLUT1 and ZO-1 (n=3 independent experiments). PCA (**e**) and correlation heatmap (**f**) showing the close clustering between biological replicates and the relative distances existing between the different samples. (**g**) Gene expression heatmap showing the genes differentially expressed either in EE-ECs vs CMEC/D3, and EE-ECs vs SM-ECs. Values are mean ± SEM of n = 3 independent experiments. ***p < 0.001. Statistical analysis is performed using Student t-test and one-way ANOVA followed by Tukey post-test. Scale bars, 100 µm. EE: ETV2, ERG1; SM: small molecules; PCA: principal component analysis.

Next, we sought to establish a reliable protocol to obtain vascular mural cells compatible with their assembly into a BBB structure starting from established procedures (*19*). Following the developmental pathway of this cell lineage, hESCs were differentiated into neural crest progenitor cells (NCPCs), as cellular intermediates, by culturing in the presence of the BMP/TGFβ inhibitors NOGGIN, SB431542, the WNT pathway activator CHIR99021 and FGF2 in N2 medium for 12 days (Fig. 2a). Subsequently, NCSCs were purified by NGFR+ immunopanning and differentiated into mural cells in N2 medium with 10% FBS for 10 days. With this procedure, a maximum of 74.6 ± 3.1% of the differentiated cells acquired the expression of the mural cell marker NG2 (Fig. 2b), indicating that a non-marginal cellular fraction failed to complete this differentiation pathway. Thus, we decided to complement this procedure with the forced expression of FOXC1, a TF essential for the developmental cell lineage specification of brain vessel-associated pericytes (*20,21*). Hence, hESC lines with the stable integration of a lentiviral vector carrying FOXC1 under the dox-inducible promoter (2.1 copies in average per cell) were established (Fig. 2c,d). F-hESC lines maintained unaltered the pluripotent stem cell state as suggested by the constitutive expression of key markers and exhibited a normal karyotype (Supplementary Fig. 1a,b). Next, F-hESC lines were differentiated using the same two-step protocol in the presence of dox for two weeks. At DIV25, nearly all cells were immunopositive for NG2 indicating a complete differentiation into mural-like cells (F-MCs). A similar result was obtained when the cell purification step was removed and cells were maintained in serum-free conditions, thereby, simplifying the overall procedure in a unique direct cell differentiation process (Fig. 2e, Supplementary Fig. 5a,b). Mural cells are known for their high contractility which control constriction and dilation dynamics of blood vessels (*22*). To determine the functional maturation of F-MCs, we exposed them to Endothelin-1 (Et-1), a well-known contracting factor for these cells (*23*). F-MCs in the presence of Et-1 drastically changed their morphology with a concomitant realignment of F-actin bundles respect to their cellular axes, suggesting that F-MCs functional response to contractile cues (Supplementary Fig. 5c). We next profiled the transcriptomes of hESC-derived MCs obtained with small molecules (SM-MCs) expressing or not FOXC1 and primary mural cells isolated from autoptic human brain tissue (pMCs), which are commercially available. Principal component analysis (Fig. 2f) and unsupervised hierarchical clustering (Fig. 2g) revealed similarity between SM-MCs and F-MCs, whereas both appeared highly divergent from pMCs. GO terms associated with vascular development and mesodermal differentiation were enriched in stem cell-derived compared to primary MCs (Supplementary Fig. 3d,e). Moreover, F-MCs show the highest correlation with primary human brain mural cells in terms of global gene expression profiling in a pairwise comparative analysis (Supplementary Fig. 3f). Considering an integrated panel of distinctive markers of mural cells (both smooth muscle cells (SMCs) and pericytes), pMCs presented a low level of all their transcripts, suggesting that these cells underwent a consistent dedifferentiation process during their in vitro expansion (Fig. 2h,i). In a direct comparison between F-MCs and SM-MCs, the former cell type presented significantly higher expression levels of these markers including CSPG4, PDGFRB, CNN1, MYH11, KCNJ8, PLXNA2, IGF2 and the endogenous FOXC1, as confirmed by independent qPCR assays (Fig. 2h,i). In contrast, SM markers of parenchymal fibroblasts such as COL1A2, VIM and FBLN2 were more enriched in SM-MCs, suggesting a partial failure to suppress the expression of alternative cell identities (Fig. 2h,i). Hence, FOXC1 expression increases the fidelity of MC differentiation and shortens the procedure by eliminating the need for cell immunoselection. Since global transcriptome revealed that F-MCs expressed markers selective for both SMCs and pericytes, to distinguish between the two mural sub-types, we performed immunofluorescence staining for TAGLN (also known as SM22), an actin-binding protein expressed exclusively by SMCs (*24*). Approximately 40% of F-MCs displayed strong TAGLN expression with low NG2 levels, indicating their differentiation toward an SMC phenotype (Supplementary Fig. 5d). By contrast, the remaining cells were TAGLN-with high NG2 expression, supporting their identity as pericytes (Supplementary Fig. 5d). These results demonstrate the successful establishment of a robust protocol for generating functional MCs with the co-presence of cells with distinguished features of both SMCs and pericytes.

**Figure 2:**
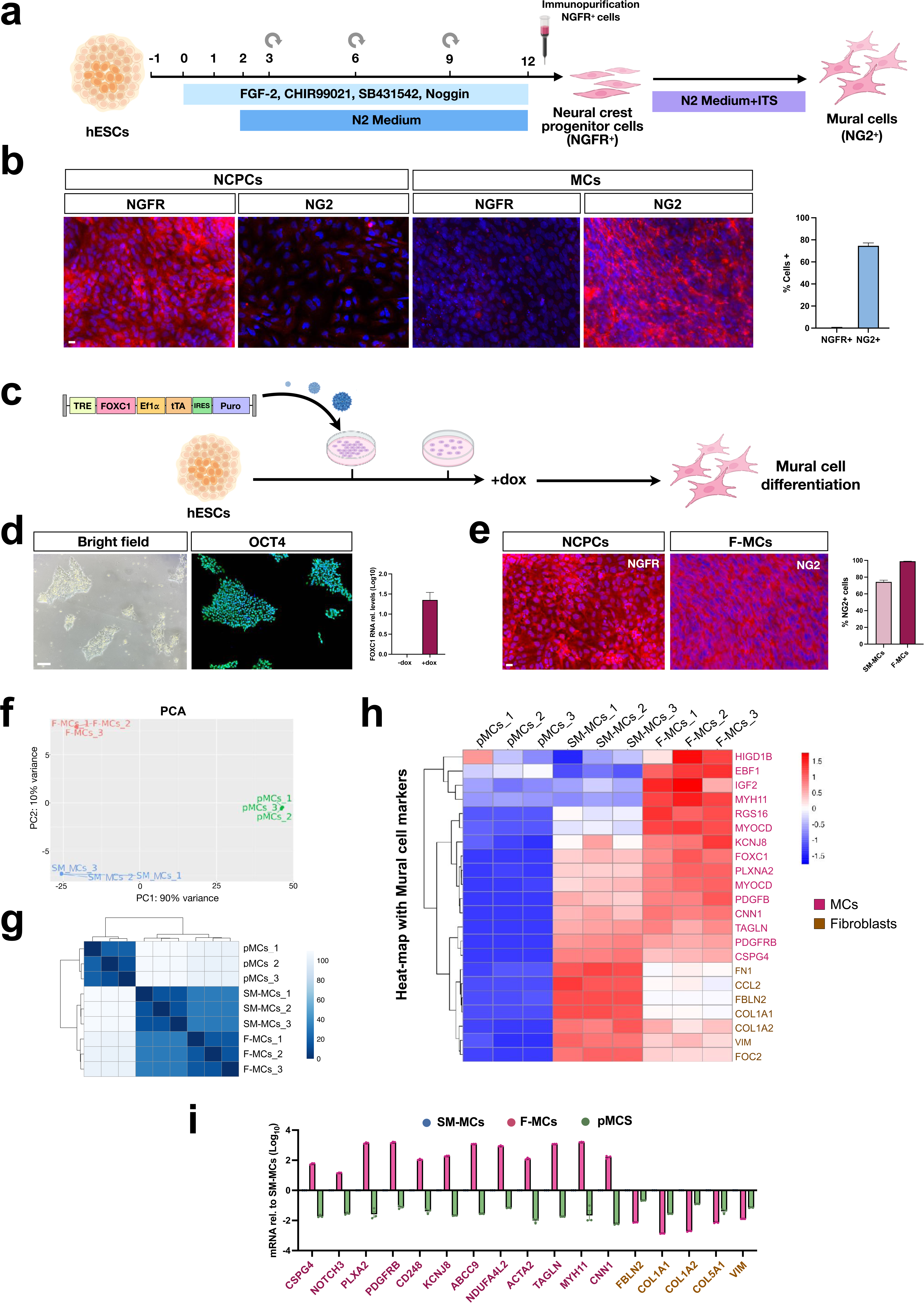
Generation of hESC-derived mural cells. (**a**) Schematic representation of the methodology for differentiating mural cells from hESCs. (**b**) Representative images and quantification of neuronal crest progenitor (NCPCs) and mural cells (MCs) immunostained for NGFR and NG2, respectively (n=3 independent experiments). (**c**) Illustration of the procedure for generating FOXC1 stable hESC lines. (**d**) Representative images of hESC colonies and qPCR analysis of FOXC1 expression with or without doxycycline (n=3 independent experiments). (**e**) Representative images and quantification of NCPCs and MCs derived from FOXC1-hESCs. PCA (**f**) and distance matrix analysis (**g**) showing the close clustering between biological replicates and the relative distances between the different samples. (**h**) Gene expression heatmap showing the genes differentially expressed either in pMCs vs SM-MCs, and pMCs vs F-MCs. (**i**) qPCR analysis confirms the differences in transcriptional levels of key marker genes between SM-MCs, F-MCs and pMCs. Quantification was normalized to SM-MCs. Values are mean ± SEM of n = 3 independent experiments. **p < 0.01, ***p < 0.001. Statistical analysis is performed using Student *t*-test. Scale bars, 100 µm. NCPCs: Neuronal crest progenitor cells; MCs: Mural cells; F: FOXC1; dox: doxycycline.

Astrocytes are the third cellular component of the BBB that we set out to obtain by differentiating hESCs with another protocol that, like the previous ones, integrates TF expression with small molecules. NFIA expression was shown to induce a gliogenic switch in hESC-derived neural progenitor cells (NPCs) promoting their differentiation into astrocytes (*25*). In the same vein, we have showed that NFIA contributes to the direct reprogramming of fibroblasts into functional astroglial cells (*26*). Therefore, we generated an hESC line with the stable integration of a lentivirus carrying a NFIA dox-inducible expression cassette (N-ESCs, with 3.2 copies/per cell) (Supplementary Fig. 2a,b). N-ESC-derived NPCs were induced into astroglial cells (N-ACs) by expansion in the AGM astrocyte growth medium (Lonza®) supplemented with dox for 7 days (Supplementary Fig. 6a,b). Three weeks after culture in this medium, nearly all NPCs were differentiated into N-ACs, 70 ± 3% of which were immunodecorated for GFAP (Supplementary Fig. 6c). qPCR assays confirmed that N-ACs expressed increased levels of markers of mature astrocytes including GLUL, SLC1A, AQP4, GFAP and VIM (Supplementary Fig. 6d). The replacement of serum with BSA in the differentiation medium did not affect N-AC differentiation as shown by comparable numbers of GFAP+ and GS+ N-ACs (Supplementary Fig. 6e). Thus, the combination of cell lineage-determining master factors together with inducing small molecules was instrumental in establishing the differentiation procedures for the efficient and rapid generation of the three BBB cellular components with high specificity and maturation state.

### Generation of mature and functional hESC-derived BBB assembloids on transwells

Next, we decided to assemble EE-ECs, F-MCs and N-ACs together on a transwell filter, where EE-ECs and F-MCs were physically plated together on the top of the filter and N-ACs were allowed to adhere to the bottom side of the filter in order to reconstitute the native BBB cellular organization (Fig. 3a). Barrier integrity was inferred by determining the transendothelial electrical resistance (TEER), a widely used quantitative technique to measure electrical resistance across a cellular monolayer. When the filter was populated with a layer of EE-ECs or CMEC/D3 cells alone, the barrier resistance was very low at 12 weeks after plating, with TEER values below 400 Ω/cm^2^ (395 and 170 Ω/cm^2^ for EE-ECs and CMEC/D3 cells, respectively) (Fig. 3b). However, the coordinated seeding of the three hESC-derived BBB cell types generated a structure that, progressively during the 12 weeks in the transwell (the final time point analyzed), reached a peak of 3480 Ω/cm² of TEER, which is within the range of values estimated for the BBB in vivo (Fig. 3b) (*27*). The increase in TEER was accompanied by a corresponding sharp reduction in permeability to sodium-fluorescein (NaFl), a hydrophilic small molecule tracer, with a 12- and 18-fold decrease of NaFl passage compared to the EC and CMEC/D3 monolayers, respectively (Fig. 3c). Next the cell-permeable fluorescent dye, Rhodamine 123 (Rho123), was used to evaluate the functional activity of the P-glycoprotein (P-gp) efflux transport system in thBBBAs (*28*). Rho123 accumulation was increased significantly in thBBBAs treated with the P-gp inhibitor cyclosporin A, suggesting the presence of functional P-gp in thBBBA endothelial cells (Fig. 3d). Rho123 is a small lipophilic dye (380 Da) which can passively diffuse into and between cells. In fact, we detected Rho123 fluorescence in the medium of the bottom compartment of the thBBBA, which increased in the presence of cyclosporin A, suggesting an effective crossing of the molecule through the barrier (Fig. 3e). Another hallmark of mature BBB is its low level of transcytosis (*29*). Consistent with this, the thBBBA showed an eightfold reduction in uptake of the transcytotic marker Tf-FITC compared to the EE-EC monolayer alone (Fig. 3f). Supporting these functional characteristics, electron microscopy revealed well-developed tight and adherens junctions, visible as dark, electron-dense lines at sites where adjacent endothelial cell membranes are closely apposed (Fig. 3g). As the stereotyped organization of EE-ECs, F-MCs and N-ACs on the transwell generated structures that exhibited the cardinal functional features of the BBB, we named them as thBBBAs for transwell-assisted human BBB assembloids. We next wondered whether changing the relative position of the F-MCs in the lower compartment in direct contact with N-ACs would have influenced the functional properties of the structure (Supplementary Fig. 7a). In fact, both longitudinal measurements of TEER and NaFl showed that this configuration (Fbottom) is significantly less impermeable (Supplementary Fig. 7b,c), suggesting that the intimate association between EE-ECs and F-MCs is an important morphological requirement for a fully functional BBB. Another critical parameter is the relative ratio between EE-ECs and F-MCs in the system. We measured TEER values in thBBBAs with varying numbers of EE-ECs while maintaining a constant number of F-MCs (40.000 cells/cm^2^) (Supplementary Fig. 7d). Notably, the increased reduction of the optimal ratio of 3:1 between EE-ECs and F-MCs was associated to a corresponding increased loss of TEER values (Supplementary Fig. 7e). This demonstrates that the relative proportion of endothelial and mural cells is essential for generating high-quality thBBBAs. To assess the cellular composition within the thBBBA, we devised a cell disaggregation treatment followed by cell sorting with selective cell surface markers (CD31+/CD49f+ for EE-ECs, CD49f+/CD31-for N-ACs and CD140b+ for F-MCs) to separate the three cellular components (Fig. 3h) (*30*). Importantly, the relative numbers between the cell-types did not significantly diverge between thBBBAs at 4 or 12 weeks of culture, suggesting the long-term stability of this cellular system in vitro (Fig. 3h). Next, isolated EE-ECs from thBBBA at 4 weeks in culture or grown in isolated monolayer for the same time were tested for targeted transcriptional analysis. Genes associated with BBB mature properties such as cell-cell junctions (SLC2A1, OCLN, TJP1, CLDN5), selective solute transport (e.g. SLC01A2, SLC38A3, SLC7A8) and reduced transcytotic vesicular trafficking (MFSD2A) were all robustly stimulated in EE-ECs from thBBBA compared to EE-ECs from isolated monolayers (Fig. 3i). Thus, the multicellular nature of the thBBBA is crucial to enhance the expression of BBB-specific components that bolster the development of its functional properties. thBBBAs are also very stable structures in vitro since functional properties (e.g. TEER values) and expression of endothelial and mural markers did not significantly change after 8 weeks in culture as assessed by global RNA-sequencing (Supplementary Fig. 8a,b).

**Figure 3:**
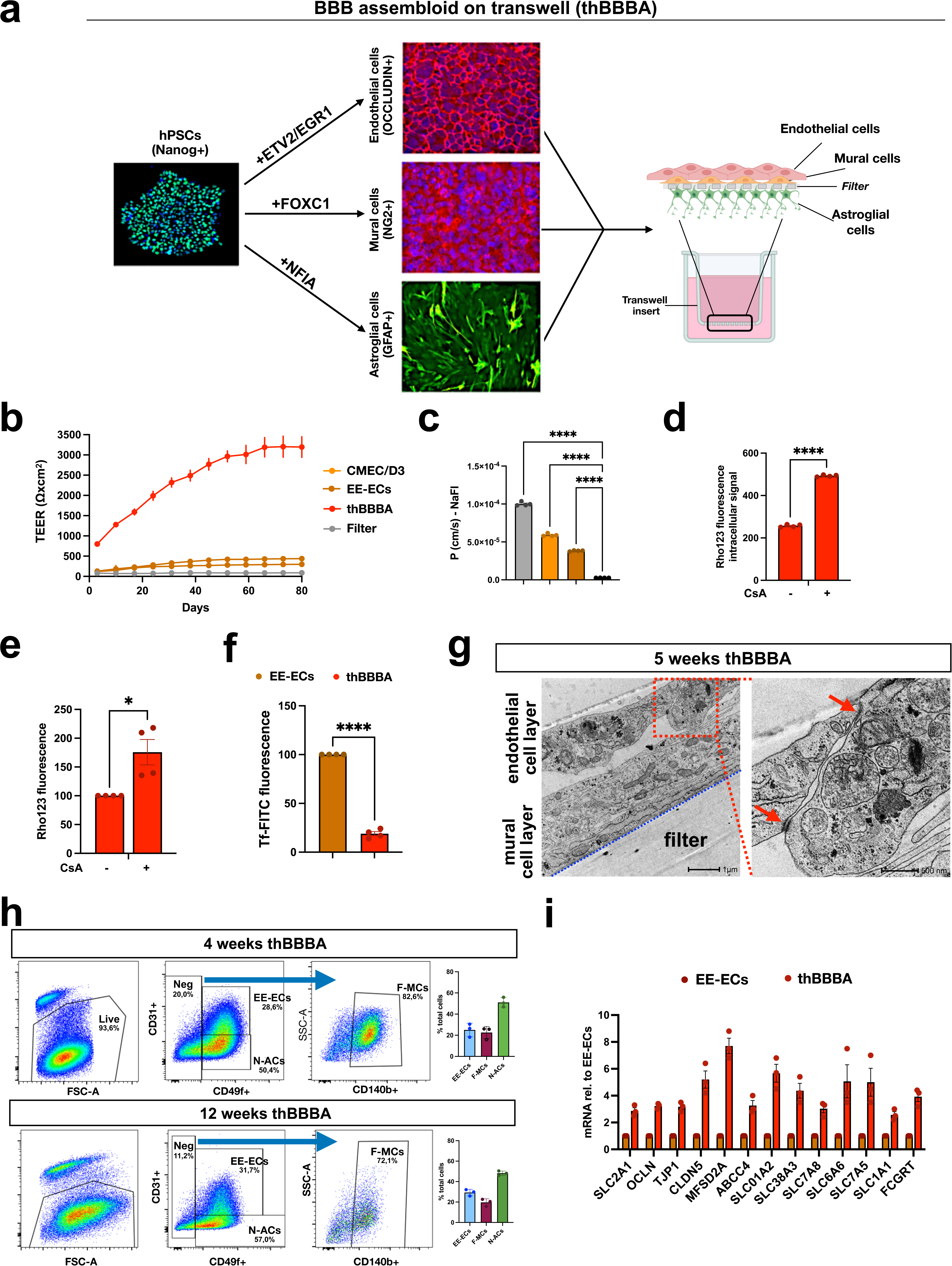
Generation of mature and functional hESC-derived BBB assembloids on transwells (thBBBAs). (**a**) Schematic view of the procedure to reconstitute the thBBBA on transwell. (**b**) Measurements of transendothelial electrical resistance (TEER) reveal that thBBBAs have significantly higher barrier resistance respect to EE-EC or CMEC/D3 cell monolayers (3 biological independent experiments). (**c**) Quantification of NaFl levels confirms the high impermeability of thBBBAs (n=3 biological independent experiments). (**d**) Quantification of P-glycoprotein transporter activity using the rhodamine 123 (Rho123) cell internalization assay with or without cyclosporin A. (**e**) Apical to basolateral flux of Rho123 in the presence or absence of cyclosporin A. (**f**) Quantitative analysis of transcytosis activity, assessed via transferrin–FITC uptake. (**g**) Representative electron microscopy images of thBBBAs with well-formed tight junctions between endothelial cells (red arrows). (**h**) Cell sorting analysis shows the relative percentage of the three cellular components in the thBBBAs at 4 and 8 weeks of in vitro culture. (**i**) qPCR analysis confirms the difference in transcriptional levels of selected key marker genes between EE-ECs and SM-ECs. Quantification was normalized to SM-ECs. Values are mean ± SEM. *p < 0.05, ****p < 0.0001. Statistical analysis is performed using one-way ANOVA followed by Tukey post-test. thBBBA: Transwell-assisted human blood brain barrier assembloids; CsA: cyclosporin A.

When starting from two unrelated iPSC lines previously generated in our lab (*31*), we obtained comparable results in terms of cell differentiation efficiencies and thBBBA functional properties, confirming that the same differentiation protocols are applicable to different PSC lines with comparable results (Supplementary Fig. 9). A key advantage of the thBBBA model is that all its cellular components are generated directly from a renewable stem cell source. Thus, to streamline the use of this cellular system, we asked whether functional thBBBAs could be generated from frozen immature intermediates of the three cell types. We therefore prepared frozen samples of immature EE-ECs, F-MCs and N-ACs and confirmed their high post-thaw viability using FACS-based analysis of live/dead double-staining (97.9% ± 2% for EE-ECs, 93 ± 4% for F-MCs and 99.2 ± 0.7% for N-ACs) (Supplementary Fig. 10b). Thawed cells were expanded for one or a few passages and then plated together on the transwell filter. thBBBAs generated with frozen or fresh cells exhibited comparable functional properties with similar TEER and sodium-fluorescein impermeability (Supplementary Fig. 10c,d). Thus, frozen cells retain the competence to generate functional thBBBAs with a simple plating step, providing an attractive plug-and-play system that can be easily adopted in any laboratory.

### Poly(I:C) enables patient NR1 auto-antibodies to cross the thBBBA

Given its fully human origin and pronounced functional properties, thBBBA is an ideal in vitro system for disease modeling. Anti-NMDA receptor (NMDAR) encephalitis is an autoimmune disorder caused by autoantibodies that target and inactivate the NR1 subunit of this receptor complex, causing severe psychiatric and neurological symptoms including hallucinations, psychosis and seizures (*32,33*). Although the disease is treatable with multiple immunotherapies, almost all patients show incomplete clinical responses and relapses are well recognized. A well-known origin of NR1-autoantibodies are ovarian teratomas that generate tertiary lymphoid germinal centers for active production of the immunoglobulins (*34*). However, peripheral autoantibodies are largely unable to cross the BBB and it is therefore unknown how they ultimately succeed in entering the brain to fuel the pathological process. This knowledge gap is also due to the lack of appropriate experimental systems that recapitulate the human-specific pathophysiological milestones of this disease. Therefore, we exploited thBBBAs to investigate the biological mechanisms leading to brain penetration of the pathogenic autoantibodies. We obtained sera and cerebrospinal fluids from three healthy donors and five patients with a definite diagnosis of anti-NMDAR encephalitis, according to the autoimmune encephalitis diagnostic criteria (Supplementary Table 1) (*35*). For a simple and sensitive readout of the successful crossing of the thBBBAs by patient autoantibodies, we seeded hPSC (either hESC or hiPSC)-derived cortical neurons that displayed NR1 immunoreactivity in the lower compartment of the transwell (Fig. 4a). Thus, patient autoantibodies bound to the NR1 subunit on the surface of hPSC-derived neurons could be visualized by anti-human IgGs fluorescence and assessed for co-localization with NR1+ puncta using a selective antibody. In fact, when patients’ sera or cerebrospinal fluids (CSFs) were diluted 1:100 in basal medium and added into the transwell, NMDAR autoantibodies were readily visualized and found to co-localize with NR1+ puncta on the human cortical neurons plated in the lower chamber (Fig. 4b). In stark contrast, the presence of the thBBBA completely prevented the crossing of the patient autoantibodies from the top to the bottom compartment, as no human IgG staining was detectable on the NR1+ puncta of the human cortical neurons (Fig. 4b). Since sporadic observations have described that the first acute signs of encephalitis can be preceded by flu-like symptoms (*36,37*), we reasoned that an infection-induced inflammatory event could contribute to a loss of BBB integrity. To investigate this possibility, we loaded the upper compartment of the transwell with patients’ serum or CSF in basal medium with or without poly(I:C) or LPS, to mimic a viral or bacterial infection, respectively (Fig. 4c). Only in the presence of poly(I:C) patients’ autoantibodies were readily detectable on human cortical neurons with 8-fold higher number of puncta compared to the challenge with LPS (Fig. 4c). Similarly, poly(I:C) was much more effective than LPS in opening the BBB, causing a marked loss of TEER values and aberrant NaFl permeability (Fig. 4c). As expected, cell junctions appeared deranged by either treatment as highlighted by the disorganized OCCLUDIN localization on the endodermal cell layer of the thBBBA (Supplementary Fig. 11a). Of note, barrier integrity was not fully rescued even 6 days after exposure to either agent, suggesting their long-term detrimental effects on the thBBBA performance (Supplementary Fig. 11b).

**Figure 4:**
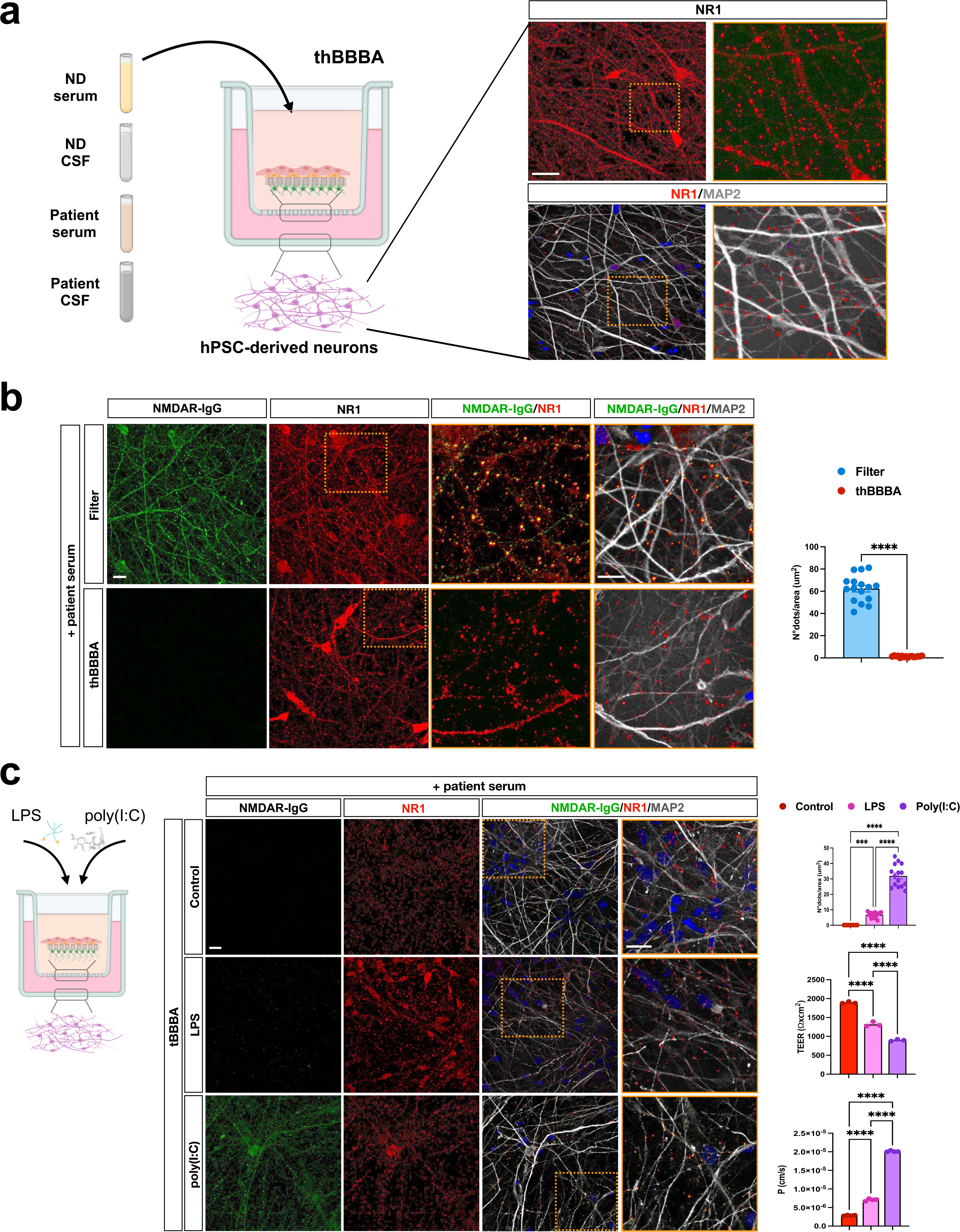
thBBBA permeability to patients’ NR1 auto-antibodies after incubation with poly(I:C). (**a**) Illustration of the anti-NMDAR autoantibodies migration assay. Immunostaining for NR1 subunits in hPSC-derived neuronal cultures plated in the bottom compartment of the transwell. (**b**) Representative images of hPSC-derived cortical neurons after treatment with the patient serum. Quantification of human IgG+ puncta in hPSC-derived neuronal cultures after exposure to the patient serum, which confirms the thBBBA high impermeability (5 fields; n=3 biological independent experiments). (**c**) Representative images and quantification of human IgG+ puncta in hPSC-derived neuronal cultures following incubation with poly(I:C) and LPS. (5 fields; n=3 biological independent experiments). Treatments with poly(I:C) and LPS cause a loss of TEER values and enhanced NaFl permeability (n=3 biological independent experiments). Values are mean ± SEM. ***p < 0.001, ****p < 0.0001. Statistical analysis is performed using one-way ANOVA followed by Tukey post-test. Scale bars, 100 µm. ND: Normal donor; CSF: Cerebrospinal Fluid; LPS: Lipopolysaccharide; poly(I:C): Polyinosinic:polycytidylic acid.

### IL-6 causes thBBBA leakage by inducing an endothelial-to-mesenchymal cell state transition which disrupts cell-cell junctions

To elucidate the causes of these adverse consequences, we measured a panel of cytokines released by the thBBBAs in the medium 48 hrs after the exposure to poly(I:C) or LPS through a bead-based immunoassay. Both compounds induced stereotyped combinations of cytokines with poly(I:C) being associated with high levels of IL-6 and TNFα in the medium, whereas LPS was associated with TNFα and IFNγ, but at much lower levels (Fig. 5a). To determine the autocrine effect of these cytokines, we exposed the thBBBA to each of these molecules at the same concentrations measured in the medium after the treatment with the immunostimulants. When patient’s serum was added to the medium supplemented with TNFα (50 ng/ml) or IFNγ (50 ng/ml) for 48 hrs, only minimal amounts of NR1 autoantibodies were able to cross the thBBBA and bind to human neurons (Supplementary Fig. 12). Conversely, an extensive human IgG staining was retrieved in human cortical neurons when the thBBBA was exposed to a medium containing IL-6 (100 ng/ml) for 48 hrs, suggesting a strong barrier leakage (Fig. 5b). In fact, IL-6 treatment caused a rapid and drastic loss of TEER values, which persisted for the following six days, confirming the significant impairment of thBBBA integrity (Fig. 5c). Inflammatory stimuli are known to promote mesenchymal features in endothelial cells, causing cytoskeletal changes, cell-cell junction rearrangements and fibroblastoid morphology. This process is referred to as endothelial-to-mesenchymal transition (EndMT) and is commonly observed to various degrees in pathologies affecting peripheral organs, but less is known about these events in BBB-specific diseases (*38,39*). Consistently, the IL-6 exposure induced key EndMT markers like SNAIL1/2, TGFβ1, TWIST and FN1 (Fig. 5d). Conversely, IL-6 downregulated key molecular components of endothelial cell junctions, including ZO-1, CDH1, and EPCAM, indicating a loss of barrier integrity (Fig. 5d). In parallel, protein levels of OCCLUDIN and ZO-1 were also reduced four days after IL-6 exposure, further confirming the disruption of the thBBBA endothelial barrier (Fig. 5e). Collectively, these results demonstrate that poly(I:C) is sufficient to open the thBBBA for NR1 autoantibodies crossing, by stimulating an autocrine IL-6 signal which promotes an EndMT process and disrupts the integrity of the thBBBA.

**Figure 5:**
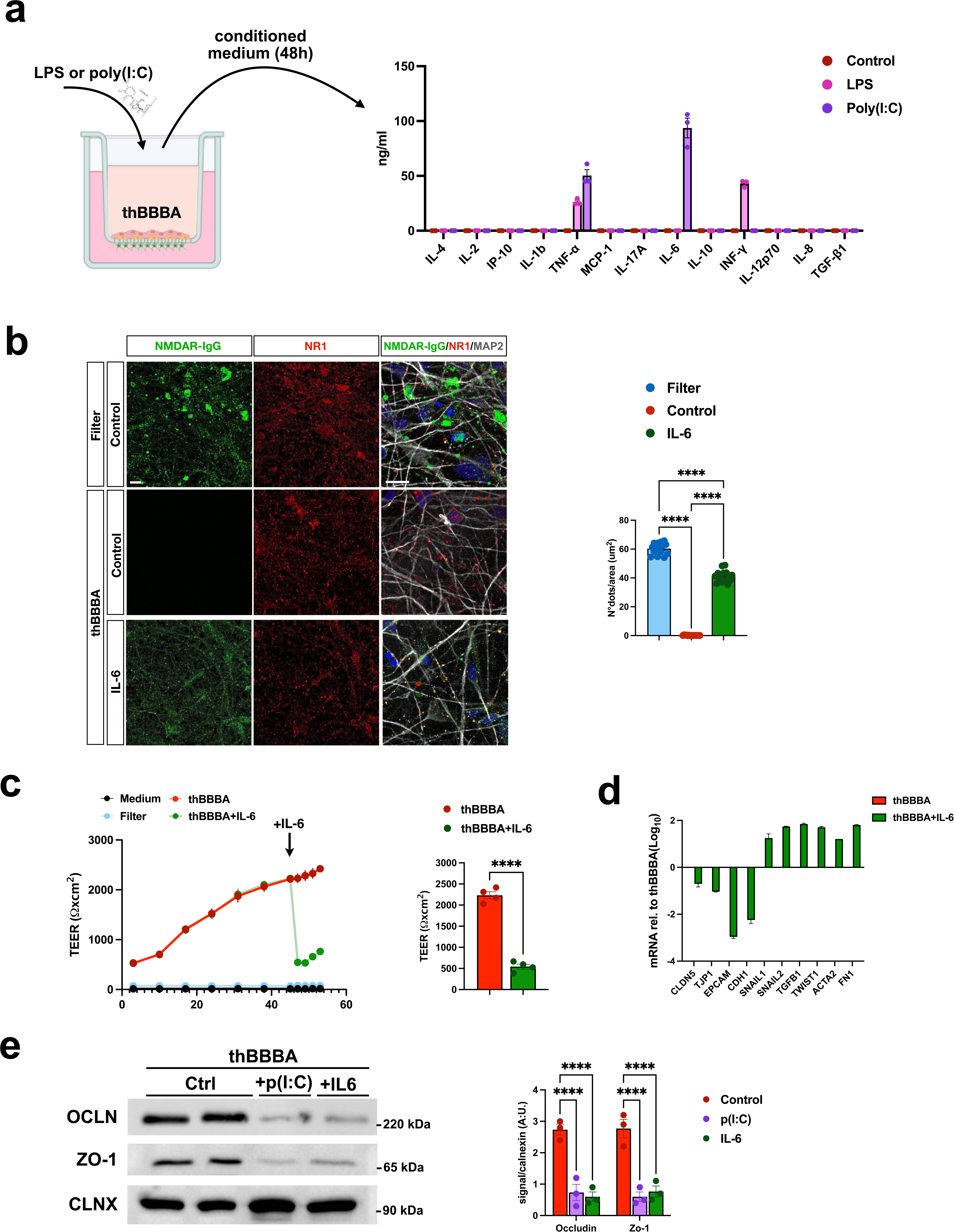
IL-6 mimics the detrimental effects caused by poly(I:C) on thBBBA integrity. (**a**) Quantification of inflammatory cytokines released by the thBBBA following incubation with poly(I:C) or LPS. (**b**) Representative images and quantification of human IgG+ puncta in hPSC-derived neurons crossing the thBBBA following incubation with IL-6 (5 fields; n=3 biological independent experiments). (**c**) Quantification of TEER levels shows a reduction in barrier integrity following incubation with IL-6 (3 biological independent experiments). (**d**) Poly(I:C) treatment induces an epithelial-to-mesenchymal transition state confirmed by the induction of key gene markers. (**e**) Immunoblot analysis and quantification reveal a reduction in tight junction markers following the treatment with p(I:C) or IL-6 compared to control. Values are mean ± SEM. ****p < 0.0001. Statistical analysis is performed using one-way ANOVA followed by Tukey post-test and two-way ANOVA followed by Bonferroni post-test. Scale bars, 100 µm.

### CAR and tocilizumab rescue poly(I:C)-induced thBBBA leakage and NR1 auto-antibody crossing

To determine the causal relationship between the IL-6-dependent EndMT process and thBBBA leakage, we though to inhibit this molecular pathway for assessing its impact on the thBBBA functional rescue. To this end, we combined the TGFβR1 inhibitor RepSox with or without the Wnt activator CHIR99021 and db-cAMP (referred to as CAR small-molecule combination). A similar group of factors has been shown to improve the barrier tightness of haematopoietic stem cell-derived endothelial cells (*40*). Poly(I:C)-induced thBBBA leakage was progressively restored to normal levels following six days of treatment with CAR compounds, as indicated by TEER and NaFl crossing measurements (Fig. 6a,b). Indeed, NR1 autoantibodies failed to cross the thBBBA in the presence of poly(I:C) together with CAR for 48 hrs (Fig. 6c). CAR mixture suppressed the activation of mesenchymal gene expression while simultaneously promoting the transcription of cell–cell junction markers, including CLDN5 (Fig. 6d). Thus, EndMT inhibition by this small molecule cocktail prevents the thBBBA permeability caused by poly(I:C). However, this treatment modulates pleiotropic and ubiquitous signaling pathways with multiple functions in different cell types, and therefore its therapeutic potential is limited, as serious side effects are expected after systemic administration. Given our data, we reasoned that blocking IL-6 signaling might be sufficient to counteract at least some of the deleterious effects of poly(I:C). To test this, we employed tocilizumab (TCZ), a humanized monoclonal antibody that acts as an IL-6 receptor antagonist. TCZ is an approved clinical product that provides substantial and long-lasting benefits in patients with rheumatoid arthritis through regular intravenous injections (*41,42*). Therefore, TCZ was added after poly(I:C) to determine its ability to rescue thBBBA dysfunction. Notably, TEER levels, NaFl impermeability, gene expression and autoantibodies crossing were all progressively restored to normal levels with continuous treatment with TCZ for up to 6 days (Fig. 6e-h). Thus, TCZ is able to restore the thBBBA tightness by inhibiting IL-6 signaling within a clinical concentration range.

**Figure 6:**
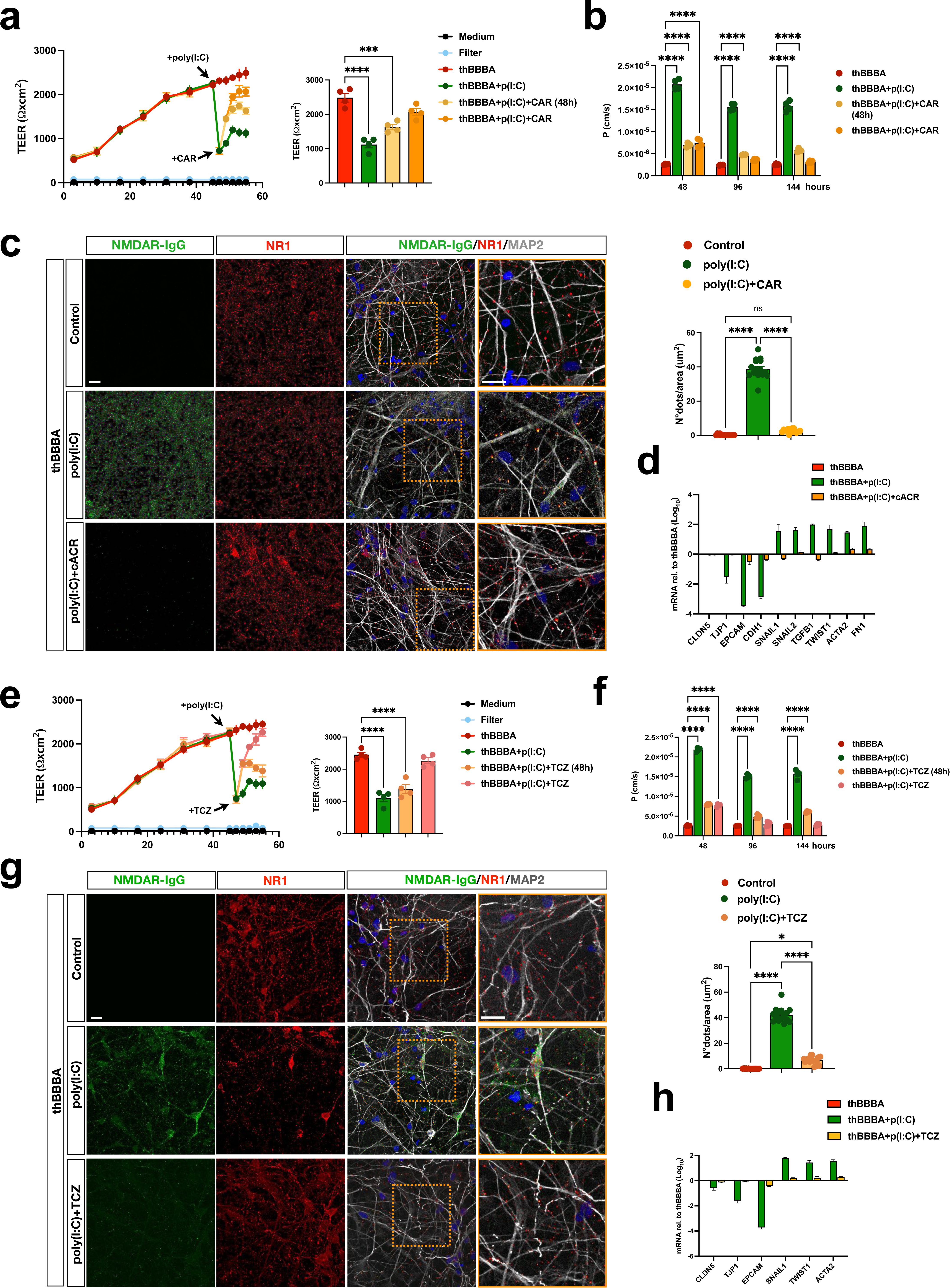
CAR and tocilizumab rescue poly(I:C)-induced thBBBA breakage and patients’ NR1 autoantibody crossing. (**a**) Quantification of TEER levels shows a reduction in barrier integrity following incubation with poly(I:C) and a recovery after treatment with the CAR factors (3 biological independent experiments). (**b**) Quantification of NaFl levels confirms the recovery after treatment with the CAR factors (n=3 biological independent experiments). (**c**) Representative images and reduction of human IgG+ puncta in hPSC-derived neurons following incubation with the CAR factors (5 fields; n=3 biological independent experiments). (**d**) CAR factors reverse the mesenchymal cell transition induced by Poly(I:C). (**e**) Rescue of TEER values after treatment with tocilizumab (TCZ) (n=3 biological independent experiments). (**f**) Quantification of NaFl levels confirms the recovery after treatment with the TCZ (n=3 biological independent experiments). (**g**) Representative images and quantification of human IgG+ puncta in hPSC-derived neuronal cultures following incubation with poly(I:C) with or without TCZ (5 fields; n=3 biological independent experiments). (**h**) TCZ treatment is sufficient to rescue the endothelial-specific gene expression as confirmed by qPCR analysis. Values are mean ± SEM. *p < 0.05, ****p < 0.0001. Statistical analysis is performed using one-way ANOVA followed by Tukey post-test. Scale bars, 100 µm. CAR: CHIR99021 (C), db-cAMP (A), RepSox (R); TCZ: Tocilizumab.

We next investigated whether a similar mechanism contributes to the pathogenetic effects of viruses known to target the brain vasculature (*43*). To this end, thBBBAs were infected with either SARS-CoV-2 (Omicron BA.1 variant) or West Nile virus (WNV) at a multiplicity of infection (MOI) of 1. Within 24 hours of inoculation, thBBBAs showed a marked reduction in TEER values, indicating substantial impairment of barrier integrity (Supplementary Fig. 13a). Cytokine profiling revealed IL-6 as the most elevated inflammatory cytokine in the culture medium, and co-treatment with TCZ partially prevented the TEER decline (Supplementary Fig. 13b,c). Altogether, these findings suggest that IL-6 signaling contributes significantly to Poly(I:C)-induced thBBBA leakage, partly mirroring mechanisms observed during infections with pathogenic viruses targeting brain endothelium.

### Ranking novel engineered AAV capsids for their capacity to cross the thBBBA or infect its endothelial cell layer

Of the natural AAV serotypes, only the AAV9 has some ability to cross the mature BBB after intravascular infusion which, however, is generally insufficient to sustain a valuable therapeutic effect (*44,45*). Recently, a growing number of engineered AAV9 capsids have been produced with enhanced brain transduction (*46,47*). However, this behaviour is generally not conserved across species, reflecting the selective binding to its target receptor in a species-specific manner. Thus, even for novel AAV capsids with high brain tropism in non-human primates (NHPs), it remains uncertain whether the same behavior is conserved into the human setting. In fact, the lack of highly reliable and predictive human-specific experimental models undermines the assessment of their clinical readiness. We thought that thBBBA might fill this gap and used it as a model to rank AAV capsids for their ability to cross a fully human BBB system and infect human neurons. To validate this application, we selected a panel of natural and engineered AAV capsids with different predicted BBB crossing abilities, in particular AAV2, which, unlike AAV9, is known to be completely unable to enter the brain tissue; and the engineered AAV9 capsids CAP-Mac, CAP-B10, AAV-Se2 and BI-hTFR1, which have been shown to have superior brain targeting in NHPs and in vitro human cells (*48–51*). A dose of 1×10E12 vg for each individual AAV capsid was added in the thBBBA medium for 6 hrs and, subsequently, the number of viral particles that successfully crossed the thBBBA and reached the bottom compartment was quantified by qPCR assays (Fig. 7a). This experiment was also replicated at 4°C, a temperature at which active transcytotic vesicle movements are suppressed and, therefore, selective transcellular viral trafficking is abolished. Under these conditions, only AAV particles that crossed the thBBBA by passive paracellular transport through cell junctions would have reached the lower compartment. As expected AAV2 was completely unable to cross the thBBBA in this setting (Fig. 7b). Conversely, all the engineered capsids showed a slightly higher penetrance compared to the unmodified AAV9. However, the BI-hTFR1 capsid showed an unmatched crossing ability with a 50-fold higher performance compared to the AAV9 (Fig. 7b). The crossing of all AAV capsids was abolished at 4°C, indicating that the successful penetration of the thBBBA followed a transcellular pathway (Fig. 7b). We next asked whether AAV particles crossing the thBBBA were able to successfully transduce human neurons. Thus, AAV infections were repeated in a thBBBA system where hPSC-derived neuronal cultures were plated into the bottom compartment (Supplementary Fig. 14a). In this setting, neuronal transduction efficiency was directly proportional to the thBBBA crossing efficiency, suggesting that infectivity of human neurons was largely comparable between the different capsids (Supplementary Fig. 14b,c). Since the superior brain transduction of these AAV capsids is not conserved in mice, this prevents a direct comparison with murine-specific AAVs, such as the PHP.eB (*52*). To allow a direct comparison between these incompatible AAVs, we generated an EE-hESC line with the stable expression of the murine Ly6a, which encodes for the PHP.eB receptor in BBB endothelial cells (Fig. 7c). Ly6A-ESCs were then differentiated into EE-ECs and assembled into the thBBBA together with F-MCs and N-ACs. As such, the Ly6A-thBBBA is suitable for infection with murine (PHP.eB) and non-murine specific AAVs (Fig. 7d). Thus, we repeated the infections on both unmodified thBBBAs and Ly6A-thBBBAs with the same protocol used before to assess a panel of AAV capsids which included the PHP.eB (Fig. 7e). As expected, the PHP.eB capsid crossed efficiently only the Ly6A-thBBBA while its penetration levels of the unmodified thBBBA were comparable to the native AAV9 (Fig. 7f). Noteworthy, only the BI-hTFR1 among the engineered AAVs achieved a thBBBA crossing efficiency close to that of the PHP.eB capsid, albeit reduced by roughly 15% (Fig. 7f). Altogether, the thBBBA, which expresses the murine receptor for the PHP.eB capsid, is a reliable system for direct comparison of BBB crossing capabilities between NHP-specific AAVs and this benchmark mouse-selective AAV. In addition to AAV9-like capsids with enhanced brain penetration, novel AAV capsid variants have been developed for improved and selective transduction of brain endothelial cells (*49,53–55*). For these AAVs as well, there is a lack of substantial evidence that their enhanced tropism is conserved in the human setting. Thus, we generated mature human EE-ESC-derived endothelial cell layers to assess and compare their transduction capabilities by measuring the amount of ZsGreen+ cells. In contrast to the mouse system, 1×10E12 AAV-BR1 viral particles infected only 15% of the human endothelial cells, similar to native AAV9 (Supplementary Fig. 15a). Conversely, same viral doses of AAV-BI30, AAV-Se1 and AAV-X1.1 showed substantially higher transduction levels, with the first capsid having the best infectivity rate with 74% of transduced cell (Supplementary Fig. 15a,b). To determine the potential to cross the BBB, these AAVs were tested in the thBBBA model. In a timeframe of 6 hrs, all 6 AAVs presented a minimal capacity to cross the thBBBA, with the AAV variants mostly unable to transverse the thBBBA in contrast with the native AAV9 (Supplementary Fig. 15c). Thus, these AAV variants, with the exception of the AAV-BR1, showed a significantly higher infectivity of the human endothelium associated with a negligible crossing capability, compared to the native AAV9.

**Figure 7:**
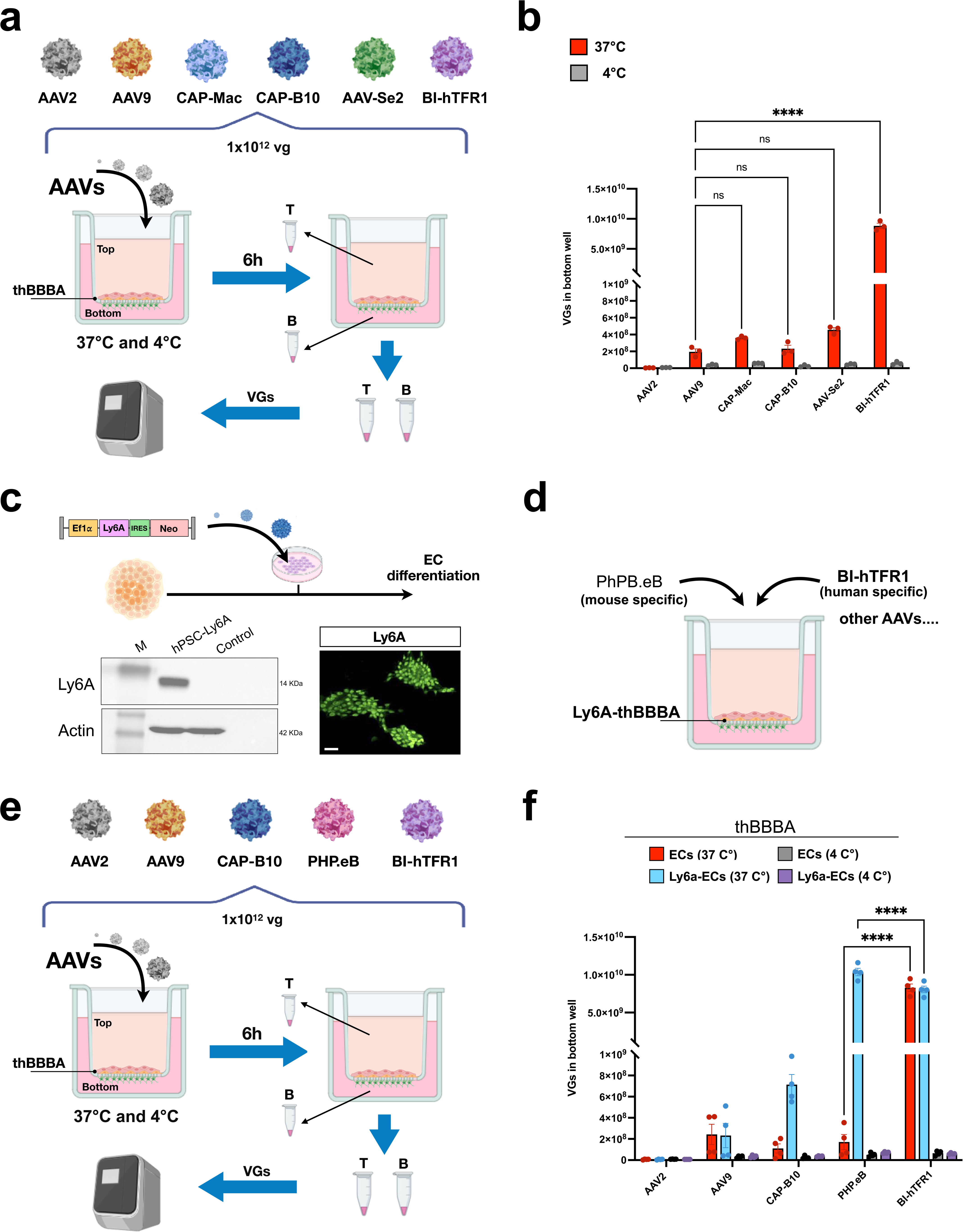
Comparative assessment of native and engineered AAVs for their capacity to cross the thBBBA. (**a**) Illustration depicting the experimental procedure used to quantify the AAVs crossing the thBBBA (**b**) qPCR quantification of the AAV vector genomes (VGs) in the lower compartment representing the AAV fraction that successfully crossed the thBBBA (n=3 biological independent experiments). (**c**) Schematic representation of the procedure for generating EE-hESC lines stably expressing Ly6a. Immunoblot and immunofluorescence confirm the expression of Ly6A in hESCs. (**d**) Schematic view of the thBBBA reconstitution with Ly6A-expressing EE-ECs. Schematic view of the experimental procedure (**e**) and quantification of VGs (**f**) showing a significant increase of the PHP.eB capsid crossing through the Ly6A-thBBBA (n=4 biological independent experiments). Values are mean ± SEM. ***p < 0.001. Statistical analysis is performed using one-way ANOVA followed by Tukey post-test and two-way ANOVA followed by Bonferroni post-test. Scale bars, 100 µm. AAV: Adeno-Associated Virus; VGs: Viral Genomes; T: Top; B: Bottom.

## Discussion

Herein, we established the thBBBA as the first model fully reconstituted by human PSC-derived cells that acquired levels of tightness and impermeability similar to the mature BBB structure in vivo. For its generation, we developed optimized protocols for endothelial and mural cell generation, combining inducing factors with the expression of cell lineage-determining TFs. These procedures ensured high differentiation efficiency and cell-type specificity of the cell derivatives. In addition, these protocols accelerated the maturation trajectory of the differentiated cells allowing, for example, ECs to form a cell layer with well-organized cell-cell junctions. In fact, the functional maturation of hPSC-derived cells is a common challenge to generate mature organ-like structures in vitro. The same issue applied for the generation of mature astroglial cells, which required the early expression of the gliogenic factor NFIA. Therefore, in this study all the three BBB cell components were derived with the aid of cell-determining TF expression, which also accelerates their functional maturation at a similar rate. The generation of fully human BBB models has previously been attempted by combining PSC-derived cells with EC lines or primary cell types isolated from autoptic human brain tissues. However, we have showed that CMEC/D3 endothelial cells have lost the expression of key molecular markers of this cell type and have failed to form a layer with high resistance and impermeability. Similarly, we found that primary human MCs displayed an extensive altered transcriptomic profile, indicating a substantial dedifferentiation state, probably induced during the culture expansion. In light of these results, the incorporation of any of these cell types into BBB models independently by their design and configuration (monolayers or multilayers in transwell or microchips) is likely to interfere with the full acquisition of their functional properties. The parallel derivation of the cellular components of the BBB from hPSCs guarantees the production of uniform and consistent cell sources for this model. Remarkably, thBBBA exhibits high impermeability with elevated TEER values. These properties arise from the maturation of endothelial-specific functional characteristics rather than from aberrant cell fate commitment, as thBBBAs do not express appreciable levels of representative epithelial genes.

These differentiation procedures yield comparable results starting from both hESCs and hiPSCs, indicating that any authentic hPSC line can be used to generate the thBBBA. Remarkably, the thBBBA is entirely derived from hPSCs, offering the unique opportunity to generate renewable sources of its cellular components. To this end, we generated precursors of EE-ECs, F-MCs and N-ACs and showed that they can establish fully functional thBBBAs after freezing. In this setting, thBBBA generation requires only a single step of cell thawing followed by the reconstitution of the model and 5 weeks of culture to achieve functional maturation. This process can be performed in any laboratory without specific expertise in hPSC biology, providing a convenient and widely accessible plug-and-play system. To determine its value, we employed the thBBBA to model anti-NMDAR encephalitis and to understand the pathogenetic mechanisms that lead peripheral autoantibodies to cross the BBB. To this end, we developed an elegant system combining the thBBBA with human hiPSC-derived cortical neuronal cultures to assess the crossing of the anti-NMDAR autoantibodies and visualize their binding to the NR1 epitope on neurons. Interestingly, we found that the poly(I:C) treatment induced a strong leakage of the thBBBA through the release of an autocrine source of IL-6. In fact, this is the most produced cytokine by the thBBBA exposed to poly(I:C), which is sufficient to disrupt the thBBBA endothelial cell junctions and allow the patients’ autoantibody to cross over. Similar effects were also obtained after infections with two viruses, SARS-CoV-2 and WNV, known to target the brain endothelium and induce pathological deficits (*56,57*). The thBBBA molecular profiling revealed an EndMT process occurring in the thBBBA endothelial cells as the principal cause of the barrier opening by the autocrine IL-6, which could be corrected by a cocktail of factors capable of inhibiting this cell state transition. We also demonstrated that TCZ protects the thBBBA from the damaging effects caused by IL-6, either produced by poly(I:C) or the neurotropic virus inoculations. TCZ is an approved clinical treatment for rheumatoid arthritis and systemic lupus erythematosus, due to the role of IL-6 in B and T cell activation (*58,59*). In addition, beneficial effects have been described in few sporadic clinical cases of refractory autoimmune encephalitis treated with TCZ (*60,61*). Our data revealed the detrimental local effects of IL-6 on the BBB and provided a rationale for the clinical use of TCZ in autoimmune encephalitis to reduce or delay the brain penetration of patients’ pathological autoantibodies. Growing evidence suggests that intrathecal antibody synthesis plays a significant role in anti-NMDAR encephalitis (*62*). However, it remains unknown how and when B or plasma cells can enter the brain to fuel the progression of the pathology. Our thBBBA system might provide a valuable setting to dissect the conditions that enable these immune cells to invade the CNS.

A limitation of the current thBBBA model is the lack of microglia cells that, although are not necessary for maintaining BBB integrity under steady-state conditions, they influence BBB behavior during pathological processes. However, the current thBBBA model has offered us the unique opportunity to define the intrinsic response of endothelial and mural cells to inflammatory stimuli without the confound effects could be played by the co-presence of microglia. Future experiments are warranted to assess the specific contribution of microglia in this system. In addition, a future thBBBA model using iPSCs generated by patients with NMDAR encephalitis might help to uncover individual-specific vulnerabilities that can facilitate the brain penetration of pathogenic autoimmune antibodies.

Given the human nature of the thBBBA, we believed that this system could fill an important gap in the validation of novel neurotropic AAVs in terms of their translational potential. In fact, many novel engineered AAV capsids have recently been selected for their superior ability to cross the BBB in mice or NHPs (*48–51*). However, it is not yet known whether these acquired properties are equally conserved in the human setting. The thBBBA provided a simple and rapid system to compare the ability to cross the human BBB between these AAV variants and native AAV9. In fact, we found that most of these engineered AAVs outperformed the AAV9, but among them, the BI-hTFR1 showed an unparalleled efficiency. One caveat of BI-hTFR1 and other AAV variants is that their BBB crossing performance is abolished in mice, probably because the sequence of their epitopes on the target cell receptors is not conserved in these animals (*51*). This shortcoming makes impossible to use these AAVs in mouse models of human diseases and to directly compare their performance with that of the PHP.eB, the most efficient capsid for crossing the murine, but not the human, BBB (*52*). To overcome this limitation, we produced thBBBAs expressing the murine PHP.eB receptor Ly6A and exposed them to PHP.eB and a panel of engineered AAV particles (*63,64*). Remarkably, only the BI-hTFR1 showed a comparable BBB crossing capacity to that of PHP.eB, albeit with 15% less efficiency. These results validate the thBBBA as an invaluable system to determine the ability of newly selected AAV variants to cross a human BBB model and to compare their performance with murine-specific AAVs such as the PHP.eB. Therefore, this model provides unique opportunity to validate the translational potential of new synthetic AAV variants and offers an unprecedented BBB model for screening AAV libraries to select novel AAV variants for their improved crossing capacity through a fully human barrier.

## Acknowledgements

We thank E. Vicenzi, M. Casucci, A. Gritti and L. Naldini for providing valuable research reagents and sharing equipment. B. Deverman is acknowledged for providing the BI-hTFR1 capsid vector. We are grateful to D. Bonanomi, J. Körbellin and all members of the Broccoli’s lab for helpful discussion. We acknowledge the FRACTAL and ALEMBIC core facilities for expert supervision in flow-cytometry and confocal imaging, respectively. This work was supported by the complementary action to the EU NRRP “D34Health” (project #PNC0000001; CUP B53C22006100001) to V.B. and the project 4-DBR (#101047099) of the HORIZON-EIC-2021-PATHFINDEROPEN call to V.B. and J.M.C.

## Author contributions

A.I. performed the experiments and analyzed data; S.G. and M.L. designed, generated and produced the lentiviral and AAV vectors; S.M. performed molecular immunoblotting and flow-cytometry experiments; M.G. developed the AAV crossing assay on thBBBAs; E.E. contributed to perform and analyze the gene expression studies; G.M. supervised the computational analysis; E.C. and N.C. performed and evaluated the impact of viruses on the thBBBA; J.M.C. contributed in designing the experiments; R.I. collected and analyzed the human tissues samples and help in the data assessment; V.B. supervised, coordinated and supported the project and wrote the paper with A.I.

## Declaration of interests

The authors declare no competing interests.

## Methods

### Cell cultures

The Genea019 embryonic stem cells (ESC) line was purchased from Myocena Inc (San Diego, CA, USA). Control iPSC lines were generated as previously described (*31*). ESC and iPSC lines were stabilized and expanded in feeder-free conditions in mTeSR1 (Stem Cell Technologies) and seeded in HESC qualified Matrigel (Corning)-coated 6-well plates.

### Molecular cloning and viral infection

ETV2, ERG or FOXC1 coding regions followed by an IRES-puromycin/blasticidin resistance cassette, were cloned into lentiviral vectors under the control of the tetracycline inducible promoter. Replication-incompetent, VSVg-coated lentiviral particles were packaged in 293T cells. To obtain a ETV2/ERG stable ESC lines, cells were transduced and cultured in mTeSR medium containing puromycin (1μg/ml, Sigma-Aldrich) or blasticidin (10μg/ml, Sigma-Aldrich). After 48 hrs single colonies were picked and seeded in hESC-qualified matrigel-coated 6-well plates. Similarly, ETV2 or ERG1 stable ESC lines were generated by selection in puromycin or blasticidin, respectively. FOXC1 stable ESC lines were generated by selection in puromycin (1μg/ml, Sigma-Aldrich).

### Endothelial differentiation protocol

Endothelial cells were generated starting from previously published protocols with appropriated optimization (*11,12*). Briefly, PSCs were dissociated in cell clusters using Accutase (Sigma-Aldrich) and seeded onto low-adhesion plates in mTeSR1 supplemented with N2 (1:200, ThermoFisher Scientific), Pen/Strept (1%, Sigma-Aldrich), human NOGGIN (0.5 μg/ml, R&D System), SB431542 (5 μM, Sigma-Aldrich), BMP4 (20 ng/ml, Miltenyi) and Y27632 (10 μM, Selleckchem) to obtain embryoid bodies. The day after medium was supplemented with ActivinA (10 ng/ml, Sigma-Aldrich) and CHIR99021 (1 μM, Miltenyi) (removed at day 10); on day 2, medium was supplemented with FGF2 (20 ng/ml, ThermoFisher Scientific) (removed at day 14). On day 4, embryoid bodies were transferred to adherent conditions on Matrigel-coated plates and medium was supplemented with 25 ng/ml VEGF-A (Miltenyi) (remained for the duration of culture); on day 7, SB431542 (Sigma-Aldrich) was added at 10 μM concentration and remained for indicated duration. At day 15 cultures were dissociated using Accutase and seeded onto Matrigel-coated plates with Human Endothelial-SFM (ThermoFisher Scientific) supplemented with B27 (1:50, ThermoFisher Scientific).

### Mural cells differentiation protocol

One day before initiating cell differentiation, hESCs maintained in mTeSR1 medium were dissociated using Accutase and seeded onto Matrigel-coated plates with mTeSR1 medium supplemented with the ROCK inhibitor Y27632 (10 µM). Differentiation of neural crest cell progenitors (NCPCs) was initiated the next day by switching to a N2 medium (DMEM/F12 basal medium supplemented with N2-1:100, ThermoFisher Scientific), Pen/Strept (1%, Sigma-Aldrich), human NOGGIN (0.5 μg/ml, R&D System), SB431542 (10 μM, Sigma-Aldrich), CHIR99021 (3 μM, Miltenyi) and FGF2 (20 ng/ml, ThermoFisher Scientific). Cells were expanded by replacing the N2-NSCF medium three times a week and passaging them when reaching 95% confluence. During passaging, cells were singularized using Accutase and replated 1:6 dilution in N2-NSCF medium. At D15 of the E6-CSFD treatment, cells were dissociated using Accutase and combined with NGFR+ microbeads (20 μl per 10^7^ cells; Miltenyi), FcR blocking reagent (20 μl per 10^7^ cells), and MACS buffer (20 μl per 10^7^ cells; 0.5% bovine serum albumin + 2 mM EDTA in sterile phosphate-buffered saline (PBS) without Ca2+/Mg2+) at 4°C for 15 min. Cells were washed in MACS buffer and resuspended in 500 μl of MACS buffer. Cells were sorted through two LS columns (Miltenyi Biotec) according to manufacturer instructions and resuspended in N2 medium supplemented with fetal bovine serum (FBS, 10%, Sigma-Aldrich) and plated onto Matrigel-coated flasks. Alternatively, the sorted NCPCs were cultured in a serum-free medium. The sorted cells were resuspended in N2 medium supplemented with Insulin-Transferrin-Selenium (ITS, 1:50 ThermoFisher Scientific) and plated onto Matrigel-coated flasks.

### Astrocytes differentiation protocol

Neural progenitor cells (NPCs) were generated as previously described with appropriated optimization (*65*). Briefly, hPSCs were dissociated in cell clusters using Accutase (Sigma-Aldrich) and seeded onto low-adhesion plates in mTeSR1 supplemented with N2 (1:200, ThermoFisher Scientific), Pen/Strept (1%, Sigma-Aldrich), human Noggin (0.5 μg/ml, R&D System), SB431542 (5 μM, Sigma-Aldrich) and Y27632 (10 μM, Selleckchem). After 10 days, embryoid bodies were seeded onto matrigel-coated plates (1:100, matrigel growth factor reduced, Corning) in DMEM/F12 (Sigma-Aldrich) supplemented with N2 (1:100), non-essential amino acids (1%, MEM NEAA, ThermoFisher Scientific) and Pen/Strept. After 10 days, rosettes were dissociated with Accutase and plated onto matrigel coated-flasks in NPC media containing DMEM/F12, N2 (1:200), B27 (1:100, ThermoFisher Scientific), Pen/Strept (1%) and FGF2 (20 ng/ml, ThermoFisher Scientific). NPC-derived astrocytes were generated as previously described with appropriated optimization (*66,67*). At differentiation day −1, 90–95% confluent NPC cultures were transduced with lentivirus expressing NFIA. The day after NPCs were washed with PBS and dissociated with Accutase and seeded in low-attachment 35-mm dishes and incubated shaking at 90 rpm for 24 h at 37 °C to obtain NPC spheres. The day after, NPC medium was replaced with DMEM-F12 supplemented with 1:100 B-27, 1:200 N-2, and 5 μM ROCK inhibitor. After 48 hrs, medium was changed to the astrocyte growth medium (AGM Bullet Kit, Lonza, #CC-3186) for 15 days, shaking at 90 rpm at 37 °C, changing medium every third day. After 2 weeks, the obtained spheres were plated onto Matrigel-coated T25 flasks in astrocyte growth medium. When cells reached ∼95% confluence, spheres were removed with a tip and adhered cells were passaged to a new flask. The culture reached astrocyte purity after the third passage. Alternatively, the astrocytes were obtained under a condition that involves the use of a serum-free medium. NPC medium was replaced with DMEM supplemented with Pen/Strept (1%, Sigma-Aldrich), ascorbic acid (200 nM, Sigma-Aldrich,), EGF (20 ng/ml, ThermoFisher Scientific), FGF2 (20 ng/ml, ThermoFisher Scientific), insulin (200 μg/ml, Sigma-Aldrich) and bovine serum albumin (BSA 100 μg/ml, Sigma-Aldrich). After 2 weeks, the obtained spheres were plated onto Matrigel-coated T25 flasks in astrocyte medium serum free composed by DMEM (Sigma-Aldrich) supplemented with Pen/Strept (1%, Sigma-Aldrich), sodium pyruvate (1%, Sigma-Aldrich) and BSA (100 μg/ml, Sigma-Aldrich).

### hPSC differentiation into cortical neuronal cultures

Neural progenitor cells (NPCs) were dissociated with Accutase and plated on matrigel-coated 6-well plates (1× 300000 cells per well) in NPC medium. Two days after, the differentiation medium containing Neurobasal (ThermoFisher Scientific), Pen/Strep (1%), B27 (1:50), with SU5402 (10 µM, Sigma-Aldrich), PD0325901 (8 µM, Sigma-Aldrich), DAPT (10 µM, Sigma-Aldrich) was added and kept for 2 days. Differentiation medium was replaced every day with a fresh one on days 1 and 2. At day 3, cells were detached by Accutase solution incubation at 37 °C for 10 min to obtain a single-cell suspension. Cells were centrifuged, counted, and seeded at a density of 55,000 cells/cm^2^ onto poly-L-lysine/laminin/fibronectin-coated plates (100 µg/ml, 2 µg/ml, 2 µg/ml, all from Sigma-Aldrich,) in neuronal maturation medium supplemented with ROCK inhibitor Y27632 (10 µM) for the first 24 h. Neuronal maturation medium was composed by Neurobasal (ThermoFisher Scientific) supplemented with B27 (1:50), 2 mM glutamine, 1% Pen/Strept, BDNF (20 ng/ml, Peprotech), ascorbic acid (100 nM, Sigma-Aldrich,), Laminin (1 μg/μl), DAPT (10 μM), dbcAMP (250 μM, Selleckchem). The culture medium was replaced the next day to remove the ROCK inhibitor, and then half of the medium was replaced with a fresh neuronal maturation medium twice a week.

### Cell seeding in transwell plates

To establish the thBBBA model, astrocytes (25 × 10³ cells/cm²) were seeded onto the bottom side of the Transwell membrane. The lower surface of the Transwell membrane was pretreated with Matrigel to promote the adhesion of the astrocytes to the membrane. After 48 hrs, mural cells were plated on the apical side (40 × 10³ cells/cm²), and after another 48 hrs, endothelial cells were seeded on top of the mural cells in different numbers (80 × 10³, 100 x 10^3^ or 120 x 10³ cells/cm²) for comparative functional assessment.

### Measurement of transepithelial/endothelial electrical resistance (TEER)

STX01 electrodes with a Millcell^®^ ERS-2 system (Millipore) were used to measure the resistance of the endothelial cell cultures in a transwell plate setup. The STX01 electrodes were placed in the transwell plate making contact between the media on the upper and the lower compartments of the plate. The resistance (U) values were measured 4 times in each transwell, including a cell-free control to account for the membrane’s resistance. The Ω values were adjusted with the surface growth area (cm^2^) of the membrane to obtain the TEER (Ω/cm^2^) values of the barrier.

### Sodium fluorescein assay

Media in the upper compartment of the transwell was replaced with 30 μM sodium fluorescein (NaFl) solution diluted in 200 μL fresh EC media. After one and two hours 200 μL were collected from the lower compartment of each transwell and was aliquoted in a 96-well plate to measure the fluorescence of NaFl flowing through the membrane with a microplate reader (Perkin Elmer VICTOR3V) using Ex (l) 485 ± 10 nm and Em (l) 530 ± 12.5 nm. The cumulative amount of compound transported into the receiver compartment was quantified at predefined time intervals. Data were plotted as a function of time, and the slope of the linear portion of the curve (dQ/dt) was calculated to determine the apparent permeability coefficient (P) using the following equation:

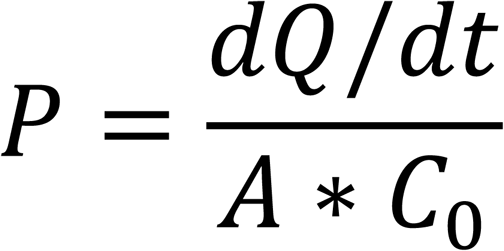

where A is the surface area of the membrane (cm²) and C₀ is the initial concentration of the compound in the donor compartment (µg/cm³).

### Efflux transporter assay

P-glycoprotein functionality was assessed using the rhodamine 123 cell internalization assay (Sigma). thBBBAs were preincubated for 1 h at 37 C with the p-glycoprotein inhibitor, cyclosporin A (5 μM, Sigma). thBBBAs were then incubated with rhodamine 123 (10 μM, Sigma) for 1 h at 37°C with or without its inhibitor. Cells were then washed three times with PBS and lysed with 5% Triton X-100 (Sigma). Fluorescence was measured using a microplate reader (Perkin Elmer VICTOR3V) using Ex (l) 485 ± 10 nm and Em (l) 530 ± 12.5 nm and normalized to cell number.

### Assessment of transport crossing the thBBBA

thBBBA were incubated with 10 μM CsA diluted only in the apical chamber for 1 h at 37°C. Next, thBBBAs were incubated with 10 μM rhodamine 123 with or without CsA for 1 h at 37°C. At the end of the incubation, 200 μL of medium was removed from the basolateral chamber, and fluorescence was measured a microplate reader (Perkin Elmer VICTOR3V) using Ex (l) 485 ± 10 nm and Em (l) 530 ± 12.5 nm Fluorescence values were normalized to medium collected from the basolateral chamber of thBBBA not treated with inhibitor.

### Transcytosis analysis

Transcytosis functionality was assessed using the Transferrin-FITC cell internalization assay (Thermo Fisher Scientific). thBBBAs were incubated with 10 mg/ml Transferrin-FITC for 20 min, then washed three times with PBS and lysed with 5% Triton X-100 (Sigma). Fluorescence was measured using a microplate reader (Perkin Elmer VICTOR3V) using Ex (l) 495 ± 10 nm and Em (l) 518 ± 12.5 nm and normalized to cell number.

### Anti-NMDAR antibodies migration assay

The upper compartment of the transwell was pretreated with the cytosolic lipopolysaccharide (LPS) (100 ng/mL, Sigma), TLR3 agonist poly(I:C) (100 ng/mL, Sigma), IL-6 (100 ng/mL, Miltenyi), TNFα (50 ng/mL, Miltenyi) or IFNγ (50 ng/mL, Miltenyi) for 48 hrs. Subsequently, the stimulus-containing medium was removed, and 200 µL of serum or cerebrospinal fluid (CSF) samples from anti-NMDAR autoimmune encephalitis patients (1:100 dilution) was added to the top compartment for 2 hrs at 37°C and immunofluorescence staining was successively performed. For the positive control, 500 µL of diluted serum or CSF was directly applied to the plated neurons. For the CAR or tocilizumab (TCZ) treatment the medium was replaced with fresh medium with dbcAMP (Selleckchem, 250 μM), RepSox (10 μM, Sigma) and CHIR99021 (10 μM, Miltenyi), (CAR factor combination) or tocilizumab (50 ng/mL, BioXCell) for 48 hrs.

### In vitro cytokines assays

thBBBAs were treated with the cytosolic lipopolysaccharide (LPS) (100 ng/mL) or poly(I:C) (100 ng/mL) for 48 hrs. Next, the co-culture supernatants were collected and analyzed using the LEGENDplex™ HU Essential Immune Response Panel (13-plex) (BioLegend). This assay is a bead-based immunoassay that uses the same basic principle as sandwich immunoassays. The beads are differentiated by size and internal fluorescence intensity, with each bead conjugated to a specific antibody for a particular analyte. When the selected panel of capture beads is mixed with a sample containing target analytes, each analyte binds to its corresponding capture beads. After washing, a biotinylated detection antibody cocktail is added, and each detection antibody binds to its specific analyte captured on the beads, forming a capture bead-analyte-detection antibody sandwich. Streptavidin-phycoerythrin (SA-PE) is then added, binding to the biotinylated detection antibodies, generating a fluorescent signal proportional to the amount of bound analyte. The fluorescent signal is measured using a flow cytometer, and the analyte concentration is determined using a standard curve. The assay FCS files were analyzed using LEGENDplex™ data analysis software, which is available together with the LEGENDplex™ assay kit.

### Live/Dead staining for viability analysis of cryopreserved cells

Cells were thawed at 37°C in culture medium and then washed with PBS. They were stained with LIVE/DEAD fixable near-IR dead cell stain kit (1:1000, Thermo Fisher Scientific) in PBS at RT for 15 minutes. After a wash in PBS, they were fixed in 1% PFA and acquired at FACS Canto II (BD). Data were analyzed with FlowJo software (Treestar).

### Separation of cellular populations in the thBBBA for Flow Cytometry

ThBBBAs were dissociated with EDTA at 37 °C for 10 minutes, collected in PBS, and centrifuged at 1000 rpm for 5 minutes. The cells were then washed with PBS and stained with antibodies against CD49f (Miltenyi, astroglial cells), CD31 (Miltenyi endothelial cells), and CD140b (Miltenyi mural cells). After an additional PBS wash, cells were fixed in 1% PFA and acquired using a FACS Canto II (BD). Data were analyzed with FlowJo software (Treestar). For sorting-based separation, cells were stained with the same antibody mix described above, washed, and sorted using a FACSAria Fusion (BD). Sorted cells were then collected and lysed in TRIzol for RNA extraction.

### Transwell assays

To test AAV transport across the thBBBA, different AAVs were added to the upper compartment of the transwell at 50,000 vg/cell (for each AAV). After six hrs of incubation at 37°C or 4°C, 50 μL of medium was extracted from the top and bottom chamber. AAVpro^©^ Titration Kit Ver2 (TaKaRa) was used to measure the number of viral particles in the two compartments. Viral lysates were treated with DNase solution for 15 minutes at 37 °C and denatured at 95°C for 10 minutes to deactivate DNase. The lysates were then diluted in 20 µL of lysis buffer and incubated in the thermocycler at 70°C for an additional 10 minutes. The fraction of AAVs that successfully crossed the thBBBAs was measured by qPCR using a standard curve using positive controls obtained from the AAVpro^©^ Titration Kit.

### Endothelin-1 stimulation of Mural Cells

Mural cells seeded on Matrigel-coated glass coverslips were exposed for 5 minutes to 100 nM Endothelin-1 (Et-1, Sigma) in HEPES/Ringer solution supplemented with 0.01% bovine serum albumin. Cells were then fixed for 20 minutes on ice with 4% paraformaldehyde (PFA, Sigma) in phosphate-buffered saline (PBS, Euroclone). After fixation, cells were washed and stained with Alexa Fluor™ 594 phalloidin (Thermo Scientific) for 30 minutes at room temperature, followed by PBS washes. Finally, the coverslips were mounted on glass slides using Dako Fluorescence Mounting Medium (Dako). Images were acquired using a Leica SP8 confocal microscope with a 63x objective.

### RNA isolation and real-time RT-PCR

RNA was extracted using the TRI Reagent isolation system (Sigma-Aldrich) according to the manufacturer’s instructions. Total RNA was treated with DNAseI (Roche) to prevent DNA contamination. One microgram of RNA was reverse transcribed using the ImProm-II Reverse Transcription System for RT-qPCR (Promega, USA). Quantitative RT-PCR (qRT-PCR) was carried out using the CFX96 Real-Time PCR Detection System (Bio-Rad, USA). One-fiftieth of the reverse-transcribed cDNA was amplified in a 16μl reaction mixture containing 1× Titan Hot Taq EvaGreen qPCR Mix (Bioatlas, Estonia) and 0.4 mM of each primer. The thermal profile consisted of 2 minutes at 50°C and 10 minutes at 95°C, followed by 40 cycles of 15 seconds each at 95°C and 1 minute at 60°C. mRNA levels were calculated according to the threshold cycle numbers within a linear range of amplification of 20 to 34 cycles. Data were standardized versus the housekeeping gene Actin, which was used as an internal standard and amplified for every sample in parallel assays.

### Electron microscopy

thBBBAs were fixed in a solution containing 2.5% glutaraldehyde and 0.5% sucrose in 0.1 M Sörensen phosphate buffer (pH 7.2) for 4–6 h. The specimens were then washed in a solution containing 1.5% sucrose in 0.1 M Sörensen phosphate buffer for a period of 6–12 h, post-fixed in 2% osmium tetroxide, dehydrated and embedded in Glauerts’ embedding mixture. Sections were cut on an Ultracut UCT ultramicrotome (Leica, Germany) and stained with uranyl acetate and lead citrate and observed on a JEM-1010 transmission electron microscope (JEOL, Japan) operating at 80 kV and equipped with a Mega-View-III digital camera and a Soft-Imaging-System (SIS, Germany) for the computerized acquisition of the images.

### Cell lysate preparation and immunoblotting

Cell lysates were prepared with a buffer containing 50 mM Tris-HCl (pH 7.4), 150 mM NaCl, 0,1% SDS and 1% Triton-X100. Protease and phosphatase inhibitor cocktails (Roche) were added immediately prior to use. The protein concentration was measured by BCA Protein Assay Kit (Thermo Fisher Scientific). Generally, 25 μg of protein lysates were loaded onto SDS-polyacrylamide gel electrophoresis and protein transferred onto a nitrocellulose membrane. The membrane was blocked with 5% no-fat dry milk in PBST (phosphate buffered saline with 0.1% Tween-20). The primary antibodies and dilutions used were the following: anti-ACTIN (1:1000, Sigma-Aldrich), anti-Ly6A (1:1000, Thermo Fisher Scientific), anti-CALNEXIN (1:2000, Sigma-Aldrich), anti-OCCLUDIN (1:1000, Thermo Fisher Scientific), anti-ZO-1 (1:1000, Thermo Fisher Scientific). Antibody incubation was followed by a horseradish peroxidase (HRP)-conjugated goat anti-mouse and anti-rabbit secondary antibodies (1:10000, Dako). The signal was revealed using the ECL-chemiluminescence kit (GE Healthcare Amersham ECL Western Blotting Detection Reagent) and detected with ChemiDoc Touch Imaging System (Biorad). Quantitation of band intensity on unsaturated exposures was performed with the Volume tool of the Image Lab 5.0 software. The adjusted values of the proteins of interest were normalized on those of Actin bands of the corresponding lanes.

### Immunocytochemistry

Cells were seeded on Matrigel-coated glass coverslips, and they were fixed for 20 min in ice in methanol or 4% paraformaldehyde (PFA, Sigma), solution in phosphate-buffered saline (PBS, Euroclone). Then, cells were permeabilized for 30 min in blocking solution, containing 0.1% Triton X-100 (Sigma-Aldrich) and 10% donkey serum (Sigma-Aldrich), and incubated overnight at 4 °C with the primary antibodies in blocking solution. Then, cells were washed with PBS and incubated for 1 h at room temperature with Hoechst and with secondary antibodies. The following primary antibodies were used: anti-OCT4 (1:100, Abcam), anti-NANOG (1:100, Abcam), anti-SOX2 (1:300, R&D System), anti TRA1-60 (1:300, Millipore), anti-CD31/PECAM1 (1:300 Bethyl Laboratories), anti-GLUT1 (1:500 Abcam), anti-OCCLUDIN (1:500 Thermo Fisher Scientific), anti-ZO-1 (1:300 Abcam), anti-NGFR (1:300 Abcam), anti-NG2 (1:500 Millipore), anti-PDGFRβ (1:300, Cell Signaling Technology), anti-αSMA (1:500, Sigma-Aldrich), anti-TAGLN/SM-22 (1:300, Abcam), anti-PAX6 (1:500 Covance), anti-NESTIN (1:300 Millipore), anti-GFAP (1:500, Agilent), anti-GLUTAMINE SYNTHETASE (GS) (1:500 Millipore), anti-NR1 (1:300 Sigma-Aldrich), anti-MAP2 (1:500, Immunological Sciences), anti-ZsGreen (1:500, Takara). Alexa Fluor^TM^ secondary antibodies were used for the immunofluorescence stainings.

### AAV production and purification

AAV replication-incompetent, recombinant viral particles were produced 293T cells, cultured in Dulbecco Modified Eagle Medium-high glucose (Sigma-Aldrich) containing 10% fetal bovine serum (Sigma-Aldrich), 1% non-essential amino acids (Gibco), 1% sodium pyruvate (Sigma-Aldrich), 1% glutamine (Sigma-Aldrich) and 1% penicillin/streptomycin (Sigma-Aldrich). Cells were split every 3-4 days using Trypsin 0.25% (Sigma-Aldrich). Replication-incompetent, recombinant viral particles were produced in 293T cells by polyethylenimine (PEI) (Polyscience) co-transfection of three different plasmids: transgene-containing plasmid, packaging plasmid for rep and cap genes and pHelper (Agilent) for the three adenoviral helper genes. The cells and supernatant were harvested at 120 hrs. Cells were lysed in hypertonic buffer (40 mM Tris, 500 mM NaCl, 2 mM MgCl2, pH=8) containing 100U/ml Salt Active Nuclease (SAN, Arcticzymes) for 1h at 37°C, whereas the viral particles present in the supernatant were concentrated by precipitation with 8% PEG8000 (Polyethylene glycol 8000, Sigma-Aldrich) and then added to supernatant for an additional incubation of 30min at 37°C. In order to clarify the lysate cellular debris were separated by centrifugation (4000g, 30min). The viral phase was isolated by iodixanol step gradient (15%, 25%, 40%, 60% Optiprep, Sigma-Aldrich) in the 40% fraction and concentrated in PBS (Phosphate Buffer Saline) with 100K cut-off concentrator (Amicon Ultra15, MERCK-Millipore). Virus titers were determined using AAVpro^©^ Titration Kit Ver2 (TaKaRa).

### RNA sequencing and bioinformatics analysis

Total RNA isolation was performed with RNeasy Mini Kit (QIAGEN). RNA libraries were generated starting from 1 µg of total RNA extracted using TRIzol (Invitrogen, Life technologies). RNA quality was assessed by using a Tape Station instrument (Agilent). To avoid over-representation of 3’ends, only high-quality RNA with an RNA integrity number (RIN) ≥ 8 was used. RNA was processed according to the TruSeq Stranded mRNA Library Prep Kit protocol. The libraries were sequenced on an Illumina HiSeq 3000 with 150bp paired-end reads using Illumina TruSeq technology. Image processing and basecall was performed using the Illumina Real Time Analysis Software. FASTQ reads were quality checked and trimmed with FastQC. High quality trimmed reads were mapped to the human genome (GRCh38/hg38) with Bowtie2 v2.3.4.3. Gene counts and differential gene expression were calculated with featureCount using latest GENCODE main annotation file and DESeq2, respectively. Geneset functional enrichment was performed with GSEA. Downstream statistics and Plot drawing were performed with R. Heatmaps were generated with GENE-E (The Broad Institute of MIT and Harvard). As in vivo benchmark, we used Seurat v5.3.0 (68) to data extracted from a snRNA human brain vascular atlas from (*16*) (Geo Accession Code: GSE163577). From their control samples, we selected cells labelled as ‘capillary’ for comparison against endothelial cells, and those labelled as ‘Pericyte’ or ‘SMCs’ for comparison against mural cells. We normalized the expression of the combined datasets for each cell type using DESeq2 v1.46.0’s Variance Stabilizing Transformation function (69). In R v.4.4.0, we assessed the similarity between pairs of samples by calculating the Spearman correlation on the transformed gene expressions. We limited this analysis to the 3000 genes with the highest variance among our bulk RNA samples.

### Virus propagation and titration

The SARS-CoV-2 Omicron BA.1 variant (hCoV-19/Italy/LOM-UniSR14/2021 GSAID Accession ID: EPI_ISL_12188061) was propagated in Vero E6 cells stably expressing TMPRSS2 (Vero E6-TMPRSS2, NIBSC 100978). Briefly, the transport medium from the nasopharyngeal swab was diluted 1:1 with serum-free DMEM supplemented with penicillin/streptomycin (P/S) and Amphotericin B and inoculated onto an 80% confluent monolayer in a T25 flask. After 1 hour of adsorption at 37°C, cells were maintained in DMEM supplemented with 2% FBS and Amphotericin B. The following day, the cell monolayer was washed and replenished with fresh medium. The cytopathic effect (CPE) was monitored using inverted phase-contrast microscopy (Olympus CKX41), and the supernatant was collected at monolayer complete disruption and stored at −80°C. West Nile virus (WNV; BEI Resources, NR-49928) was propagated in Vero E6 cells (ATCC CRL-1586). Cells were infected at ∼80% confluency in DMEM supplemented with 2% FBS and maintained at 37 °C in 5% CO₂. Supernatants were collected when CPE became evident and stored at −80 °C. All virus stocks were titrated by endpoint dilution assay (TCID₅₀/mL) in either Vero E6 or MDCK cells, depending on the virus. Briefly, cells at 95% confluency were infected with 10-fold serial dilutions of virus for 1 hour at 37°C, washed with PBS, and overlaid with complete medium. After 72 h, CPE was evaluated, and titers were calculated using the Reed-Muench method.

### Infection of thBBBAs with viral preps

*In vitro* thBBBA transwell cultures were infected with WNV or SARS-CoV-2 at a multiplicity of infection (MOI) of 1. Viral inoculum was added to the apical chamber in serum-free medium and incubated for 1 h at 37 °C to allow adsorption. Following infection, the inoculum was removed, cells were washed with PBS, and fresh complete medium was added to both the apical and the basal compartment. Mock-infected cultures, treated with medium only, served as negative controls. Transendothelial electrical resistance (TEER) was measured using an Epithelial Volt-Ohm Meter Millicell ERS-2 (Merck Millipore) equipped with Millicell ERS Probes (Merck Millipore). Measurements were performed at baseline (pre-infection) and at 3, 6, 24, and 48 hrs post-infection. After virus exposure, supernatants from the apical chamber were collected, heat-inactivated at 56°C for 30 minutes and stored at −80 °C until use for cytokine quantification.

### Statistics

All values are expressed as mean ± SEM. Differences between means were analysed using the Student *t*-test, one-way or two-way analysis of variance (ANOVA) depending on the number of groups and variables in each experiment. Data were then submitted to Tukey post-hoc test using GraphPad Prism software. The null hypothesis was rejected when P-value was < 0.05.

## Supplementary Material

**Supplementary Figure 1: Co-expression of ETV2 and ERG1 enhances endothelial marker localization in hESC-Derived Cells.** Representative images of endothelial cells derived from hESCs expressing ETV2, ERG1, or both (EE-hESCs), immunostained for PECAM1 (green), OCCLUDIN (red), and ZO-1 (red) (n=3 biological independent experiments). Scale bar, 100 µm.

**Supplementary Figure 2: Characterization of N-, EE-, F-hESCs.** (**a**) hESC colonies immunostained with the pluripotency markers OCT4, NANOG, SOX2 and Tra-1-60 and their morphology appearance in bright field. (**b**) qPCR analysis of pluripotency genes in ESCs compared to human fibroblasts (n=3 biological independent experiments). (**c**) Karyotype analysis confirmed the absence of noticeable chromosomal abnormalities in EE-, F-, and N-hESCs. Scale bars, 100 µm.

**Supplementary Figure 3: Bioinformatics analysis of bulk RNA-Seq in hESC-derived endothelial and mural cells.** (**a**) Volcano plot of differentially expressed genes in EE-ECs vs CMEC/D3, EE-ECs vs SM-ECs and EE-ECs vs SM-ECs. (**b**) Gene ontology analysis of differentially expressed genes. (**c**) Correlation indexes of gene expression pattern between native endothelial cells (GEO: GSE163577) and EE-ECs, SM-ECs and CMEC/D3 comparing the most variable 3000 genes between our cell populations. (**d**) Volcano plot of differentially expressed genes in F-MCs vs pMCs, SM-MCs vs pMCs and SM-MCs vs F-MCs. (**e**) Gene ontology analysis of differentially expressed genes. (**f**) Correlation indexes of gene expression pattern between native mural cells (GEO: GSE163577) and F-MCs, SM-MCs and pMCs, comparing the most variable 3000 genes between our cell populations. DEGs: Differentially expressed genes; GO: Gene Ontology.

**Supplementary Figure 4: CHIR99021 modulates brain-specific endothelial junctional marker expression.** (**a**) Representative images of hESC-derived endothelial cells cultured with or without CHIR99021, immunostained for PECAM1 (green), OCCLUDIN (red), and ZO-1 (red) (n = 3 independent experiments). (**b**) qPCR analysis confirms the difference in transcriptional levels of selected key marker genes between EE-ECs and EE-ECs w/o CHIR. Quantification was normalized to EE-ECs w/o CHIR (n=3 independent experiments). Values are mean ± SEM. Scale bar: 100 µm.

**Supplementary Figure 5: In vitro differentiation of hESCs into F-MCs** (**a**) F-MCs maintained in medium with serum or ITS (Insulin+Transferrin+Selenium) have similar proliferations rates. (**b**) F-MC differentiation efficiency is equivalent in medium supplemented with FBS or ITS (n=3 biological independent experiments). (**c**) Representative images of F-MCs exhibiting a pronounced reconfiguration of the actin cytoskeleton after exposure to Endothelin-1 (Et-1). (**d**) Representative images and quantification of mural cells immunostained for NG2 (red) and TAGLN (green) (n=3 independent experiments). Values are mean ± SEM. Scale bar: 100 µm.

**Supplementary Figure 6: In vitro differentiation of hESCs into N-ACs.** (**a**) Illustration of the protocol for differentiating astroglial cells from hESCs. (**b**) Representative image and quantification of neuronal progenitor cells (NPCs) derived from N-hESCs (n=3 biological independent experiments). (**c**) Representative image and quantification of NPC-derived astroglial cells immunostained with the specific markers Glutamine Synthetase (GS, red) and GFAP (green) (n=3 biological independent experiments). (**d**) qPCR analysis confirms the transcriptional upregulation of selected gene markers for mature astroglial cells between N-NPCs and N-ACs. Quantification was normalized to N-NPCs. (**e**) N-AC differentiation efficiency is equivalent in medium supplemented with FBS or BSA (n=3 biological independent experiments). Values are mean ± SEM. Scale bars, 100 µm. FBS: fetal bovine serum; ITS: Insulin (I), transferrin (T), and selenium (S); NPCs: neuronal progenitor cells; BSA: bovine serum albumin.

**Supplementary Figure 7: F-bottom thBBBA configuration.** (**a**) Schematic view of the thBBBA-Fbottom cellular organization with F-MCs plated on the lower side of the filter. (**b, c**) Measurements of TEER (**b**) and quantification of NaFl levels in thBBBA compared to thBBBA-Fbottom (**c**) (n=3 biological independent experiments). **(d, e)** thBBBA illustration with TEER values of thBBBAs related to increasing endothelial cell numbers (80.000, 100.000 or 120.000 cells/cm^2^) (n=3 biological independent experiments).

**Supplementary Figure 8: Expression analysis of BBBA cell types at 8 weeks of in vitro culture.** (**a**) Average GSVA scores compared between endothelial and mural samples and their corresponding 8 W samples (8 weeks of in vitro culture). Relevant gene sets extracted from MSigDB cell type signature gene set database. (**b**) Heatmap of normalized gene counts showing endothelial and mural key marker gene expression in the different cell samples. Values scaled per gene.

**Supplementary Figure 9: Characterization of hiPSC-derived thBBBAs.** (**a**) hiPSC colonies immunostained with the pluripotency marker OCT4 and visualized by bright field imaging. (**b**) Representative images and quantification of hiPSC-derived EE-ECs stained with the specific markers OCCLUDINS (red), GLUT1(green) and ZO-1 (red) (n=3 biological independent experiments). (**c**) Representative images and quantification of hiPSC-derived F-MCs stained with NG2 (red) and TAGLN (green) (n=3 biological independent experiments). (**d**) Representative images and quantification of astroglial cells (N-ACs) derived from hiPSCs (n=3 biological independent experiments). (**e**) Illustration of the cellular organization of the hiPSC-derived thBBBAs in the transwell. (**f**) High TEER values of the thBBBAs corroborate the generation of thBBBAs with mature barrier integrity (n=3 biological independent experiments). Values are mean ± SEM. ****p < 0.0001. Statistical analysis is performed using one-way ANOVA followed by Tukey post-test. Scale bars, 100 µm.

**Supplementary Figure 10: Generation of mature thBBBAs starting from frozen cell samples.** (**a**) Schematic outline and representative images of the key cellular components (endothelial, mural and astroglial cells) that reconstitute the thBBBA derived from their frozen precursors. (**b**) Quantification by FACS analysis of post-thaw viability using live/dead double-staining (n=3 biological independent experiments). (**c**) TEER values show no difference between conventional thBBBAs and thBBBAs derived from frozen cells (Fr-thBBBA) (n=3 biological independent experiments). (**d**) NaFl permeability test confirming comparable tightness between thBBBAs and Fr-thBBBAs (n=3 biological independent experiments). Values are mean ± SEM. ****p < 0.0001. Statistical analysis is performed using one-way ANOVA followed by Tukey post-test. ECPs: Endothelial progenitors.

**Supplementary Figure 11: Sustained barrier dysfunctions after p(I:C) exposure.** (**a**) Representative images of thBBBAs immunostained for OCCLUDIN show disrupted cell junction structure following exposure to LPS and p(I:C). (**b**) Quantification of TEER levels demonstrates reduced barrier integrity after incubation with LPS and p(I:C), which was not fully restored even 6 days after exposure to either agent (n=3 biological independent experiments). Values are mean ± SEM. **p < 0.01 ****p < 0.0001. Statistical analysis is performed using one-way ANOVA followed by Tukey post-test. Scale bars, 100 µm.

**Supplementary Figure 12: TNFα and IFNγ have minor effects on thBBBA integrity.** Representative images (**a**) and quantification (**b**) of human IgG+ puncta in hPSC-derived neurons after thBBBA treatment with TNFα (50 ng/ml) or IFNγ (50 ng/ml) (5 fields; n=3 biological independent experiments). Values are mean ± SEM. ****p < 0.0001. Statistical analysis is performed using one-way ANOVA followed by Tukey post-test. Scale bars, 100 µm.

**Supplementary Figure 13:** (**a**) Quantification of TEER levels shows a reduction in barrier integrity following infection with either SARS-CoV-2 or West Nile virus (WNV) (3 biological independent experiments). (**b**) Quantification of inflammatory cytokines released by the thBBBA infected with SARS-CoV-2 or WNV (n=3 biological independent experiments). (**c**) Rescue of TEER values after treatment with tocilizumab (TCZ) (n=3 biological independent experiments). Values are mean ± SEM. ***p < 0.001. ****p < 0.0001. Statistical analysis is performed using one-way ANOVA followed by Tukey post-test.

**Supplementary Figure 14: Neuronal transduction efficiency of AAV capsids after successful thBBBA crossing.** (**a**) The cartoon illustrates the experimental model used to quantify the viral neuronal transduction efficiency after successful thBBBA crossing. (**b-c**) Representative images (**b**) and quantification (**c**) of transduced hPSC-derived cortical neurons co-immunostained for ZsGreen and MAP2. (n=3 biological independent experiments). Values are mean ± SEM. ***p < 0.001. Statistical analysis is performed using one-way ANOVA followed by Tukey post-test. Scale bars, 100 µm.

**Supplementary Figure 15: Assessment of the transduction efficiency of brain endothelial-selected AAV capsids in EE-EC monolayers** (**a-b**) Representative images (**a**) and quantification (**b**) of AAV transduced hESC-derived EE-ECs in endothelial monolayers stained with ZsGreen (n=3 biological independent experiments). (**c**) Quantification of the AAV particles able to successfully cross the thBBBA (n=3 biological independent experiments). Values are mean ± SEM. ***p < 0.001. Statistical analysis is performed using one-way ANOVA followed by Tukey post-test. Scale bars, 100 µm.

**Supplementary Table 1:** Clinical and instrumental assessments of the NMDAR autoimmune encephalitis patients and description of their biological samples used in this study.

## Notes

### Competing Interest Statement

The authors have declared no competing interest.

